# A Sap Peptide Conserved across Flowering Plants Positively Regulates Lignin Biosynthesis, Biomass and Immunity

**DOI:** 10.1101/2024.05.20.594799

**Authors:** Chang-Hung Chen, Pin-Chien Liou, Yi-Fan Hsu, I-Fan Wang, Chun-Yu Kuo, Kuan-Hao Huang, Jhong-He Yu, Chin-Wen Chen, Chia-Chen Wu, Da-Gin Lin, Cheng-Bin Li, Yuan-Kai Tu, Chuan-Chih Hsu, Jung-Chen Su, Kai Xia, Isheng Jason Tsai, Ying-Chung Jimmy Lin, Ying-Lan Chen

## Abstract

Signaling peptides act as hormones to deliver short- or long-distance intercellular signals to govern complex developmental processes. Identifying endogenous signaling peptides is challenging due to their low abundance and the unknown cleavage sites required for release from precursor proteins, not to mention the investigation of their evolutionary roles across species. Consequently, very few peptides were evolutionarily characterized *in vivo*, especially long-distance signaling peptides. Here we present current largest peptidomic datasets from six species (maize, camphor tree, tomato, rose gum, soybean and poplar), totaling 12,242 peptides, selected from all representative evolutionary clades of angiosperms, including monocots, magnoliids, rosid eudicots, and asterid eudicots. A sap peptide was found to be identical across all six species and named as ASAP (<u>a</u>ngiosperm <u>sap</u> <u>p</u>eptide), emerging as the most conserved peptide family discovered thus far. ASAP rapidly induces a series of protein phosphorylation involved in a signaling cascade previously reported to regulate lignin biosynthesis, plant growth and plant immunity. Functional assays on ASAP activity demonstrated its capability on the induction of monolignol biosynthesis and lignin deposition. High-throughput phenomic analyses showed that ASAP significantly increased plant above- and below-ground biomass. In addition, ASAP treatment enhanced plant immunity and reduced the number of galls and egg masses against nematode invasion. This study provides insights into the conservation and functional significance of plant long-distance mobile signaling peptides, offering potential applications in crop improvement and disease management strategies.

## Introduction

Peptides can act as upstream signals to regulate various plant physiological processes^1–5^, and recently have been considered as hormones in plants^6^. The information delivered by functional peptides basically involves four steps, (1) the expression of peptide precursor genes, (2) proteolytic cleavage of precursor proteins, (3) peptide migration and (4) peptide perception by receptors^7–10^. The identification of signaling peptides have been considered challenging due to two reasons. First, because these peptides act as upstream signals, their endogenous abundance is usually extremely low. Second, the endogenous peptides with the unknown enzymatic cleavage rules usually possess more random matches than the fixed enzymatic digested peptides during database searching. Thus, currently only around 15 signaling peptide families have been identified in plants^3, 11–14^.

Similar to the structural consistency observed in other phytohormones across different plant species, the sequences of several peptide families were reported to be evolutionary conserved. Such conservation of peptide sequences was characterized by two layers of evidence, identification by *in silico* analyses using DNA sequences or by *in vivo* detection using LC-MS/MS. Take CLAVATA3 (CLV3)/EMBRYO SURROUNDING REGION (CLE) family for example, CLV3 was identified *in vivo* in *Arabidopsis thaliana*^15^, and the conservation level was extended to angiosperms and even land plants using *in silico* analyses^16, 17^. Since the generation of peptides requires enzymatic activity of protease, *in silico* identification of peptides would be risky. However, currently reported conservation of peptides in most previous studies is typically inferred from *in vivo* identification of the peptide in one species, followed by *in silico* analyses in other plants.

Signaling peptides can be roughly divided into two categories as short- and long-distance mobile peptides based on the transporting distance. Short-distance mobile peptides travel between neighboring cells to exert intra- or inter-tissue communications by symplastic or apoplastic pathways^18, 19^. For long-distance transportation, the peptides are secreted to vascular bundle and exclusively delivered by vascular sap^20, 21^. In addition to intra- and inter-tissue communications, long-distance mobile peptides can further conduct inter-organ crosstalk to synchronize the development in entire plants^22–24^. Most of current identified peptides were reported to exert short-distance signaling among neighboring cells/tissues. To date, only five peptides from three peptide families were confirmed as functional peptides capable of long-distance signaling^14, 25–28^.

Considering the difficulties in identifying novel peptide families, the challenges in characterizing cross-species peptide conservation *in vivo*, and the limited understanding of long-distance peptide signaling, new information contributed to any of the three aspects would provide fundamental insights into peptide signaling. In this study, we performed cross-species peptidomic analyses on vascular sap from six angiosperms, exemplifying all representative evolutionary clades. Using our recently established pipeline for endogenous peptide identification with liquid chromatography tandem mass spectrometry (LC-MS/MS)^2, 29^, we identified one novel peptide in vascular sap across all six species *in vivo*, which signifies the discovery of a new long-distance peptide family function. The subsequent functional assays demonstrated that this peptide regulates monolignol biosynthesis, lignin deposition, plant biomass, and plant immunity.

## Results

We performed comprehensive peptidomic analyses in vascular sap of six species across angiosperms to obtain comprehensive knowledge toward three critical aspects: the identification of novel peptide family, characterization of peptide conservation *in vivo*, and long-distance peptide signaling. These six species were selected from angiosperms, including *Zea mays* (maize) (poales, monocots), *Cinnamomum kanehirae* (camphor tree) (laurales, magnoliids), *Solanum lycopersicum* (tomato) (solanales, asterid eudicots), *Eucalyptus grandis* (rose gum) (myrtales, rosid eudicots), *Glycine max* (soybean) (fabales, rosid eudicots), and *Populus trichocarpa* (poplar) (malpighiales, rosid eudicots) (Fig. 1A).

**Fig. 1.**
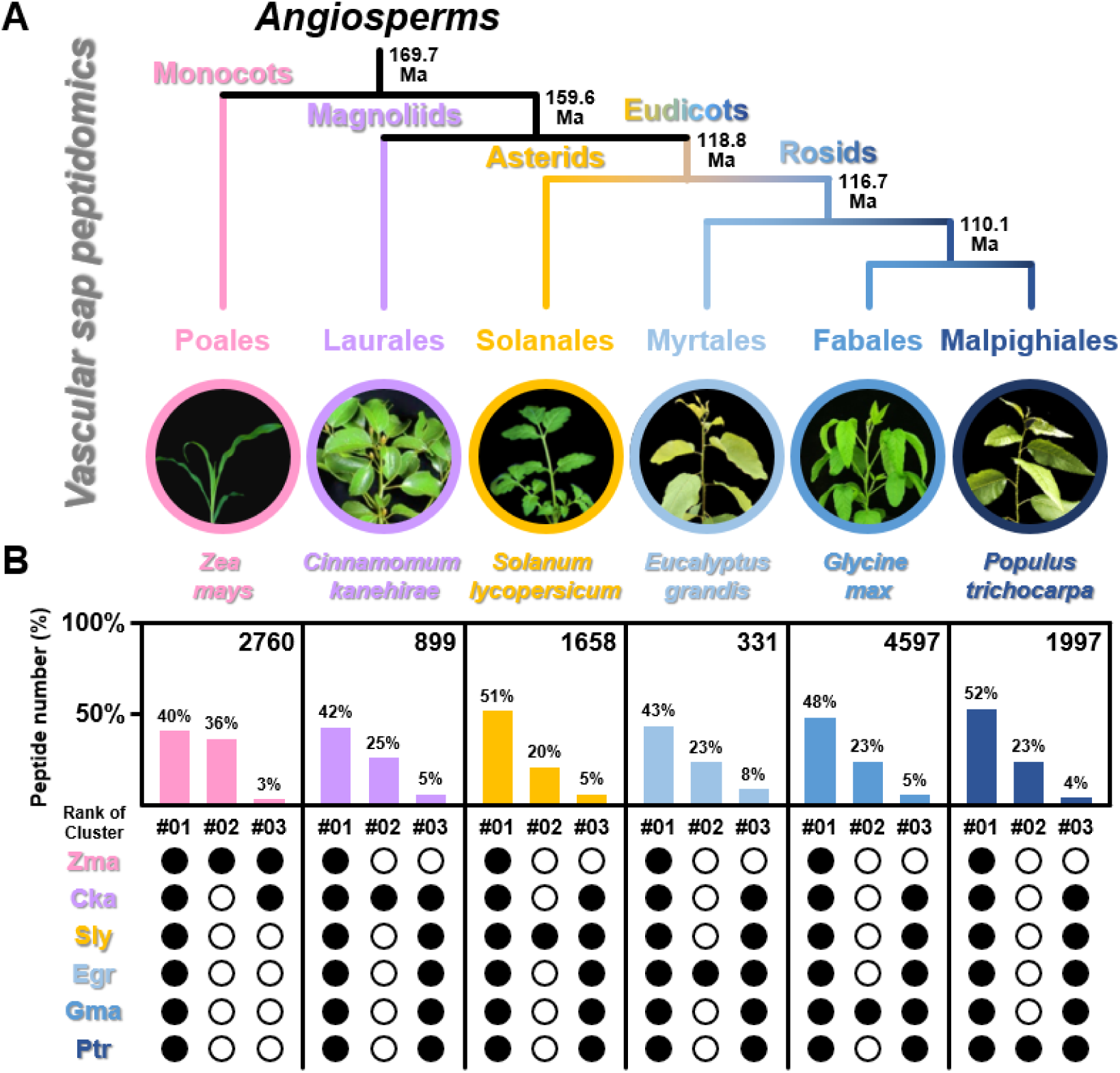
Conservation analysis for vascular sap peptidomes of six species across angiosperms. (A) Phylogeny of six species selected from different clades in angiosperms^98^, including *Zea mays* (maize) (poales, monocots), *Cinnamomum kanehirae* (camphor tree) (laurales, magnoliids), *Solanum lycopersicum* (tomato) (solanales, asterid eudicots), *Eucalyptus grandis* (myrtales, rosid eudicots), *Glycine max* (soybean) (fabales, rosid eudicots), and *Populus trichocarpa* (poplar) (malpighiales, rosid eudicots). (B) UpSet plots exhibiting the sequence conservation of total identified peptides among six species. The peptide shared ≥ 65% sequence identity in a species were considered as conserved, and the corresponding species was labeled as filled circle.

Previous studies have reported three of the largest vascular sap peptide datasets in soybean, *Medicago*, and poplar, each containing approximately 100 to 200 peptides^21, 30, 31^. With our recently established peptidomic profiling pipeline^2, 29^, we observed a significant increase in peptide numbers across all datasets from the selected six species, with an average of around 2000 peptides (Table S1). This represents a substantial fold change, with the highest dataset containing nearly 23 times more peptides than the previous largest datasets (4597/200) (Fig. 1B).

Comparative peptidomic analyses were conducted to examine the conserved and diverged peptides among these six species. Take *Z. mays* for example, the sequences of the identified 2760 peptides in *Z. mays* were used for BLAST searches to identify conserved sequences in other five species. Our recent studies have demonstrated that the CAP-derived peptide (CAPE) motif (CNYD.PxGNxxxxxPY) shares a sequence identity of 60% between *S. lycopersicum* and *A. thaliana*, suggesting a conserved function in enhancing SA-related immunity against pathogen infections^2, 32^. To identify functionally conserved peptides across angiosperms, we have raised the stringency criteria, considering peptides as conserved only if they share at least 65% identity in two species. All six peptide datasets displayed a consistent trend where the largest and second-largest populations represented two opposite cases: peptides conserved across all angiosperms and species-specific peptides, respectively (Fig. 1B; Table S2). These two populations together accounted for at least 66% of the entire dataset. Take *Z. mays* as an example, a total of 40% of the 2760 peptides were found to have homologous sequences in the other five species, while 36% of them were specific to *Z. mays* (Fig. 1B; Table S2A). Although the third-largest population only represents 3 to 8%, its conservation trend follows the evolutionary relationship: peptides from monocots are similar only to those from magnoliids, which are considered the closest evolutionary clade to monocots. Peptides in magnoliids and eudicots exhibit similarities to each other, consistent with their evolutionary relationship where magnoliids and eudicots share a common ancestor that diverged from monocots (Fig. 1B). The six largest datasets of vascular sap peptides revealed the significant roles of the peptides, either as conserved across all angiosperms or specific only to their respective species.

Amongst the conserved sap peptides, only one sequence XXXXXXXXXXX was found to be identical in all six species (Table S3). To further confirm the validity of this peptide, we extracted the MS/MS spectrum information from the datasets of each species, and the MS/MS fragments (the b and y ions) perfectly matched to each other and the synthetic control (Fig. 2A). The position weight matrix analysis using 114 species selected from 50 families in distinct clades of angiosperms showed that the peptide sequence and its flanking sequences (potential recognition site of protease) are highly conserved across angiosperms (Fig. 2B; Table S4). This identical sap peptide detected in all representative evolutionary clades of angiosperms suggested a critical biological role of this peptide on long-distance signaling, and we then named this peptide ASAP as <u>a</u>ngiosperm <u>sap</u> <u>p</u>eptide.

**Fig. 2.**
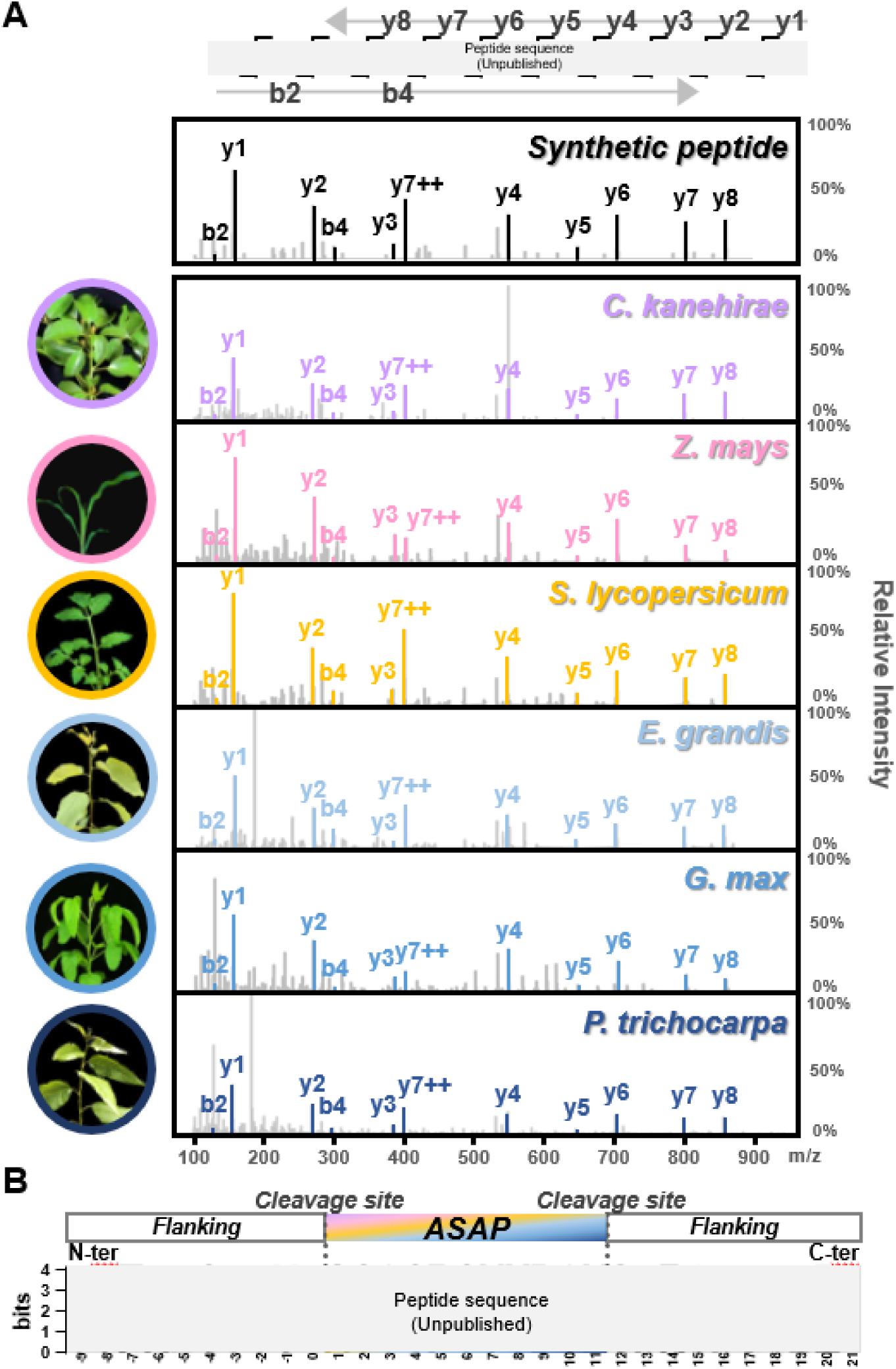
Identification of a sap peptide conserved across angiosperms. (A) Detection of endogenous ASAP (<u>A</u>ngiosperm <u>sap</u> <u>p</u>eptide) from six species (*Zea mays*, *Cinnamomum kanehirae*, *Solanum lycopersicum*, *Eucalyptus grandis*, *Glycine max*, and *Populus trichocarpa*) and synthetic ASAP using LC-MS/MS. Each spectrum exhibited the b and y fragment ions of ASAP after MS/MS fragmentation. m/z, mass-to-charge ratio. (B) Position weight matrix of aligned ASAP sequence and its 10 flanking residues extracted from total 114 species from 50 distinct families in angiosperms.

We applied two methods on sap collection to avoid the potential bias from each of the method. In first method, the stem was cut, the sap accumulated on the tree stumps were collected by pipetting (Fig. S1A). The second method centrifuged stem fragments in falcon tubes and collected sap (Fig. S1B). The first method created much less damaged sites, and the second method required much shorter processing time. We applied both methods to collect vascular sap in *P. trichocarpa* (rosid eudicots) and *S. lycopersicum* (asterid eudicots), and ASAP was successfully detected in both species using both methods (Fig. S1).

To investigate the function of ASAP, we utilized synthetic ASAP for plant irrigation to mimic the effect of ASAP overexpression. We first ensure that the synthetic ASAP could effectively enter the vascular sap through irrigation, thus mimicking endogenous ASAP. Isotope-labeled ASAP, XXXXXX[X(^13^C_5_, ^15^N_1_)]XXXX, was employed for irrigating treatment on *P. trichocarpa* and served as a standard for LC-MS/MS detection. Our successful detection of isotope-labeled ASAP in the vascular sap confirms the efficacy of the treatment (Fig. 3A). Since the majority of vascular sap is transported by mature xylem (dead xylem), the differentiating xylem (living xylem) appears to be one of the closest tissues to mature xylem, which may be a potential destination of ASAP. We then also successfully detected isotope-labeled ASAP in differentiating xylem (Fig. 3A). The detection of isotope-labeled ASAP in vascular sap and differentiating xylem demonstrated ASAP as a mobile sap peptide with the ability to enter target tissues.

**Fig. 3.**
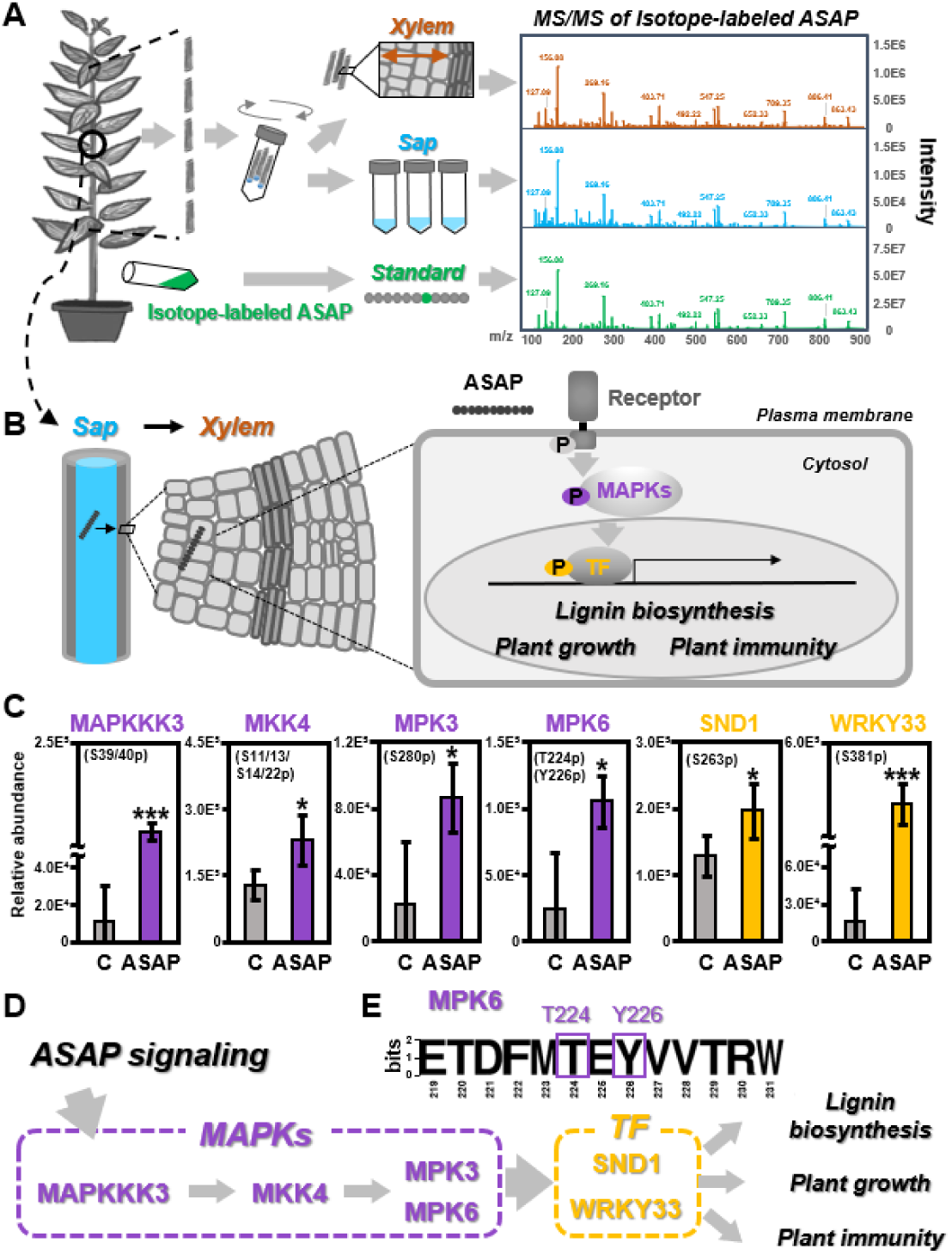
The mobility and signaling activity of ASAP. (A) Detection of exogenous isotope-labeled ASAP in sap and xylem of *Populus trichocarpa*. Synthetic isotope-labeled ASAP (labeled as Standard) was prepared as 400 pmol/injection for LC-MS/MS analysis. Peptides were extracted from the vascular sap (labeled as Sap) and stem-differentiating xylem (labeled as Xylem) that were collected from the stem tissues of the same plant after isotope-labeled ASAP treatment. The MS/MS fragments of isotope-labeled ASAP were analyzed by LC-MS/MS targeting on isotope-labeled ASAP (m/z 552.7874). The b and y fragment ions of ASAP generated by MS/MS fragmentation were labeled in all spectra. m/z, mass-to-charge ratio. (B) Proposed model of ASAP transport and signaling to regulate potential functions based on protein phosphorylation cascades. MAPKs, mitogen-activated protein kinases; TF, transcription factors. (C) The phosphopeptide levels of selected MAPKs and TFs regulated by ASAP. The debarked stems of *P. trichocarpa* treated by ddH2O (labeled as C) or ASAP were used for phosphoproteomic analyses. (D) Proposed ASAP signaling pathway via the phosphorylation of MAPKs and TF for regulating lignin biosynthesis, plant growth and plant immunity. (E) Position weight matrix of MPK6 phosphosite regions extracted from 6 species, including *Zea mays*, *Cinnamomum kanehirae*, *Solanum lycopersicum*, *Eucalyptus grandis*, *Glycine max* and *P. trichocarpa*. MAPKKK3, MAPK kinase kinase 3. MKK4, MAPK kinase 4. MPK3, mitogen-activated protein kinase 3. MPK6, mitogen-activated protein kinase 6. SND1, secondary wall-associated NAC domain 1. One and three asterisks represent Student’s *t*-test p < 0.05 and 0.001, respectively. Three individual plants were performed as biological replicates for each treatment.

Upon the perception of peptides by receptors, the most common phenomenon would be the phosphorylation of receptors and their downstream regulatory proteins, such as MAPK (mitogen-activated protein kinases) or transcription factors. We then carried out time-course phosphoproteomic analyses after ASAP treatment for 0.5, 1, 2, and 4 hours in *P. trichocarpa* (Table S5A). Debarked stems were chosen for the analyses because they contain mature xylem (where ASAP is conducted), as well as differentiating xylem and pith—the two tissues surrounding mature xylem from outside and inside. The use of debarked stem offered an ideal sample for monitoring ASAP activity. After ASAP treatment, the protein phosphorylation of many MAPK-related kinases and transcription factors within a regulatory cascade reported previously were significantly modified (Fig. 3B), including MAPKKK3^33^, MKK4^33–39^, MPK3^33, 34, 40^, MPK6^33, 34, 36–40^, NST (NAC SECONDARY WALL THICKENING PROMOTING FACTOR)^41^ and WRKY33^40^ (Fig. 3C; Table S5B). Within this cascade, MAPKKK3 located as the most upstream factor passing the signals through a series of phosphorylation on the rest of the kinases and finally to two master regulators NST3/SND1 (secondary wall-associated NAC domain 1)^42, 43^ and WRKY33^44^ for the regulation of three critical biological processes: lignin biosynthesis, plant growth and plant immunity (Fig. 3D). Among the proteins in this regulatory cascade, dual phosphorylation sites (TxY motif) on MPK6 were demonstrated to be highly conserved across plant and animal kingdoms^45^, and MPK6 thus serves as a hub for the signaling amplification. In our case, we also found two phosphorylation sites on MPK6 conserved across all six species (Fig. 3E, Table S5C), suggesting a conserved signaling pathway initiated by ASAP treatment in angiosperms.

We also observed other proteins reported to be critical members for peptide signaling (Fig. S2A and B; Table S5B). In addition to the phosphorylation of receptors, MAPKs, and transcription factors, peptide signaling usually involves other protein phosphorylation. Phosphorylated proteases would generate another type of peptides to pass the signal to the next level^3^, and phosphorylated H^+^-ATPase pumps change the pH of apoplastic environment to affect signaling^46, 47^. SIRK1 (sucrose-induced receptor kinase 1) and RLK902 (receptor-like kinase 902) receptors were reported to be involved in plant development and immunity, respectively^48, 49^. The receptor interacting protein PBL27 (PBS1-like 27) participated also in plant immunity^50^. Metacaspase 4 (MC4) is a protease for producing plant elicitor peptide 1 (Pep1)^51^, a signaling peptide that regulates lignin biosynthesis and plant immunity in *A. thaliana*^52, 53^. PEPR2 is known to be responsible for Pep1-regulated lignin biosynthesis that physically interacts with a proton pump AUTOINHIBITED H^+^-ATPase 2 (AHA2) for fine-tuning the apoplastic pH^54^, such pH changes also influence the Pep1-PEPRs interaction^47^. The phosphorylation sites of these proteins are conserved among 5 to 6 species in the selected angiosperms (Fig. S2C, Table S5C). Thus, the phosphoproteomic results indicated three potential signaling pathways induced by ASAP: lignin biosynthesis, plant growth and plant immunity, prompting subsequent functional assays to examine ASAP activities.

Lignin is one of the most critical factors for plant terrestrialization to provide mechanical support of plant body and water impermeability of vasculature. To validate whether ASAP could regulate lignin biosynthesis, the relative abundance of ten representative compounds in monolignol synthesis pathway was quantified in *P. trichocarpa* (rosid eudicots), *E. grandis* (rosid eudicots) and *S. lycopersicum* (asterid eudicots) using LC-MS/MS. In *P. trichocarpa*, two compounds participated in S-lignin pathway were significantly up-regulated, including sinapaldehyde and sinapyl alcohol (Fig. 4A and Fig. S3A). Instead, the compounds in H-lignin and G-lignin pathways exhibited no significant difference (Fig. 4A and Fig. S3A). The analyses in *E. grandis* showed highly consistent results that the metabolites toward S-lignin pathway were enriched (Fig. 4B and Fig. S3B). Although the monolignol biosynthesis pathway in *S. lycopersicum* did not show a similar trend as that in the two rosids after ASAP treatment, many metabolites in both leaves and stems were also significantly up-regulated (Fig. 4C and Fig. S3C and D). Our results demonstrated that ASAP regulates monolignol biosynthesis.

**Fig. 4.**
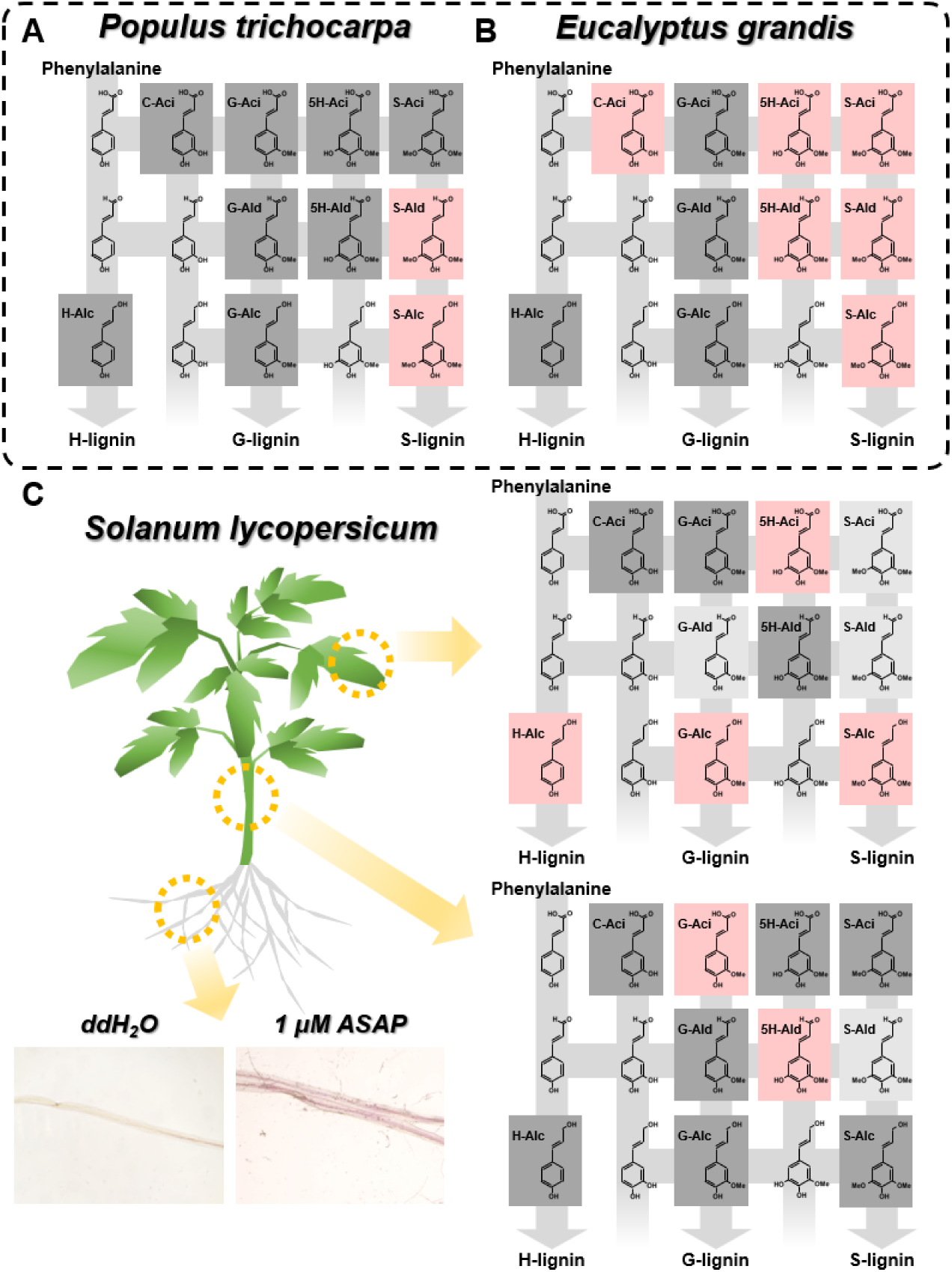
Lignin and monolignol biosynthesis positively regulated by ASAP. The changes of monolignol metabolites of (A) the debarked stems on *Populus trichocarpa* and (B) the stems on *Eucalyptus grandis* after one month of ASAP treatment. (C) Lignin deposition on roots and the changes of monolignol metabolites on stems and leaves of *Solanum lycopersicum* (tomato) after 24 hours of ASAP treatment. Ten monolignol metabolites were quantified using LC-SRM-MS analysis. Metabolites in the main fluxes of H-lignin, G-lignin and S-lignin biosynthesis are represented. The detected compounds were highlighted, pink for significant up-regulation, dark grey for no significant change, light grey for no detection. C-Aci, caffeic acid; G-Aci, ferulic acid; 5H-Aci, 5-hydroxyferulic acid; S-Aci, sinapic acid; G-Ald, coniferaldehyde; 5H-Ald, 5-hydroxyconiferaldehyde; S-Ald, sinapaldehyde; H-Alc, 4-coumaryl alcohol; G-Alc, coniferyl alcohol; S-Alc, sinapyl alcohol. Lignin staining using 5% phloroglucinol in ethanol for tomato roots. The lignified tissues were showed as the pink-red coloration. Three individual plants were performed as biological replicates for each treatment.

Lignin biosynthesis involves in monolignol biosynthesis and lignin deposition. We then tested lignin deposition through root staining of *S. lycopersicum* (asterid eudicots) under ASAP treatment (Fig. 4C). Given that the identical sequence of ASAP can be identified in *A. thaliana* protein database and that root staining of *A. thaliana* is easily observable, we thus also used *A. thaliana* (rosid eudicots) for the assessment of lignin deposition. Water treatment was used as a negative control. The treatment of a pathogen-derived peptide, flg22, was also used as a negative control due to its incapability of the induction of lignin deposition in vasculature^53^. Comparing to water or flg22 treatment, the lignin deposition was apparently enhanced upon ASAP treatment in *S. lycopersicum* and *A. thaliana* (Fig. 4C; Fig. S4A and B). The regulation of both monolignol biosynthesis and lignin deposition by ASAP demonstrates its capability in signaling for lignin biosynthesis.

Next, we examined another potential ASAP function, the regulation on plant growth. Manual quantification of plant growth, such as biomass, usually suffered from the bias generated by variation among samples and subjective judgement. To obtain objective and reliable results on plant growth, we exerted high-throughput phenomic analyses on plants using 5-15 biological replicates for *P. trichocarpa* (rosid eudicots), *S. lycopersicum* (asterid eudicots), and *Z. mays* (monocots). To avoid environmental and positional variation, we placed these biological replicates in an alternating pattern in the greenhouse (Fig. 5A and B). Digital biomass on each plant was calculated by the multiplication of plant height by leaf area (Fig. 5C), and these parameters were recorded for 14 to 28 days after water (control) or ASAP treatment (Fig. 5D to F; Table S6 to S8). All three species exhibited significant enhancements in digital biomass after ASAP treatment, with *P. trichocarpa* showing the most pronounced effect, reaching up to a 4.4-fold increase (Fig. 5D). Besides aboveground biomass, we also quantified the root development of *S. lycopersicum* by high-resolution image scanning. Total root length, taproot length, lateral root length, and root volume were all significantly increased by ASAP (Fig. S5). As a result, ASAP treatment significantly induced plant growth in all tested species across angiosperms.

**Fig. 5.**
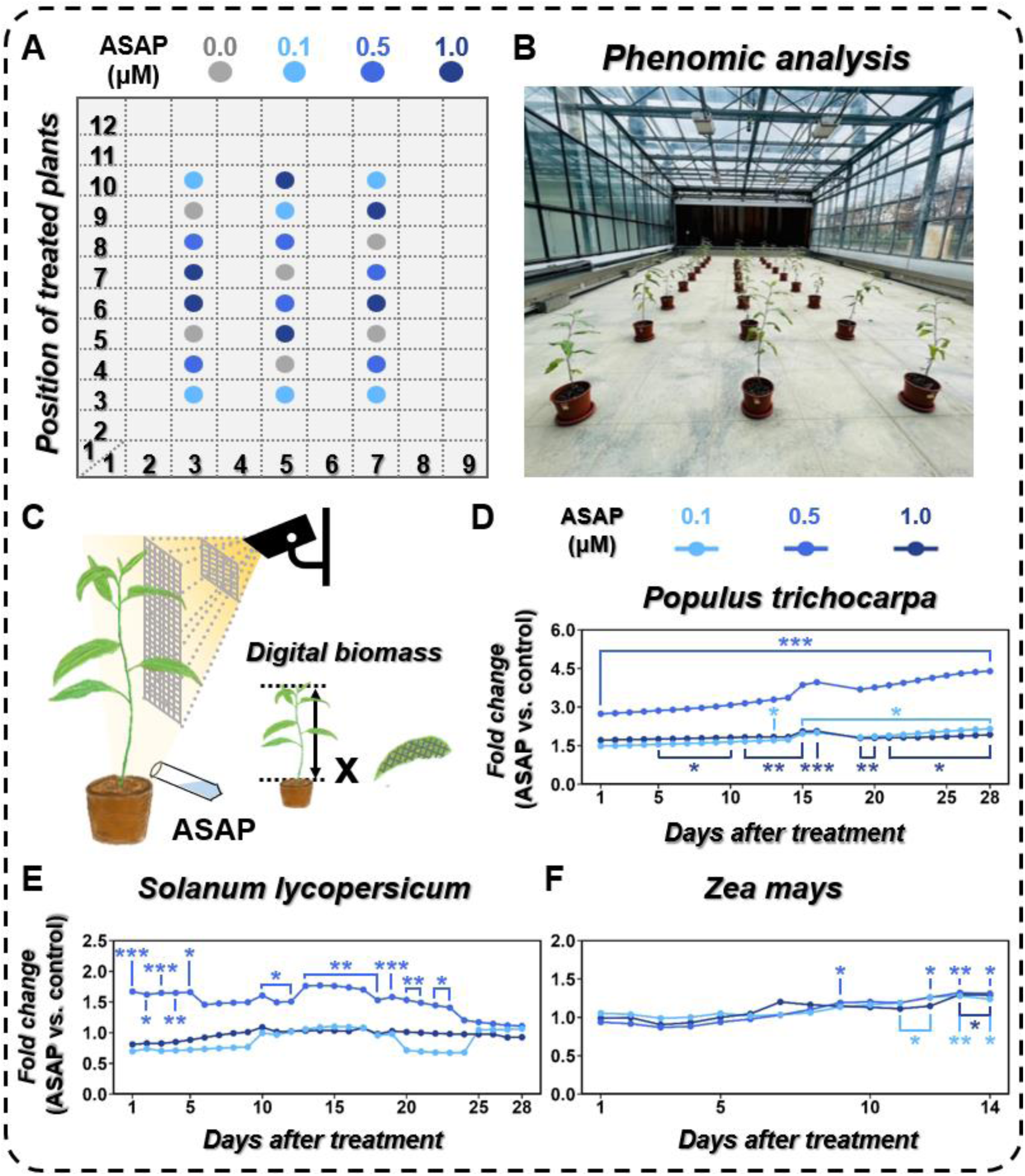
Digital biomass positively regulated by ASAP. (A) Schematic diagram shows the position arrangement of plants treated with different concentrations (0.0, 0.1, 0.5 and 1 μM) of ASAP solution in the phenotyping greenhouse. (B) Photograph of the *Populus trichocarpa* plants used for phenomic analysis in the phenotyping greenhouse. (C) Phenomic analysis for digital biomass using a high-throughput scanner, PlantEye F500 (Phenospex). The calculation of digital biomass is the multiplication of plant height by leaf area. The ASAP-induced changes of digital biomass in compared to water control (labeled as 0.0 μM ASAP in Fig 5A) in different species, including (D) *Populus trichocarpa* (n = 6), (E) *Solanum lycopersicum* (n = 5), and (F) *Zea mays* (n = 15) (see the details of calculation in Methods and Fig. S10). Each plant was irrigated with 100 ml ASAP solution or water (control) three times per week. One, two and three asterisks represent Student’s *t*-test p < 0.05, 0.01 and 0.001, respectively.

Considering the ASAP activity on *S. lycopersicum* in terms of induction on lignin biosynthesis and plant growth on both aboveground/belowground, we next investigate whether ASAP could alter plant immunity in *S. lycopersicum*. Root-knot nematodes pose a significant threat as biotrophic pests in agriculture, resulting in annual losses estimated at about $80∼100 billion^55, 56^. We inoculated root-knot nematodes by pouring around 500 larvae at J2 stage to the soil of each plant, and the vascular sap of control (without inoculation) and inoculated plants were collected (Fig. 6A). The abundance of ASAP significantly reduced after nematode inoculation (Fig. 6B; Table S9), suggesting an antagonistic relationship between ASAP and nematode development. We then examined whether ASAP treatment could enhance plant immunity against nematodes. In addition to water treatment control, we also used Pep6 peptide as control, because Pep6 was reported to involve in the regulation of plant immunity^57^. The plants were respectively treated by water, ASAP or Pep6 for 24 hours followed by nematode inoculation. ASAP significantly suppressed the number of nematode egg masses and galls. Pep6, instead, did not promote plant immunity in tomato against nematode (Fig. 6C). To explore more detailed mechanisms for the increased plant immunity triggered by ASAP, we detected the abundance on the proteins involved in the signaling of two defense-related hormones, salicylic acid (SA) and jasmonic acid (JA). SA was reported as the hormones against biotrophic pests, including plant parasitic nematodes. In contrast, JA promote wound regeneration upon nematode infection, which prompts nematode development. Comparing to water treatment, ASAP significantly induced the abundance of SA responsive protein (Pathogenesis-related protein 1, PR1 in Fig. 6D), and reduced JA biosynthesis protein (lipoxygenase, LOX in Fig. 6D). Therefore, ASAP enhanced plant immunity through the regulation of hormone signaling pathways.

**Fig. 6.**
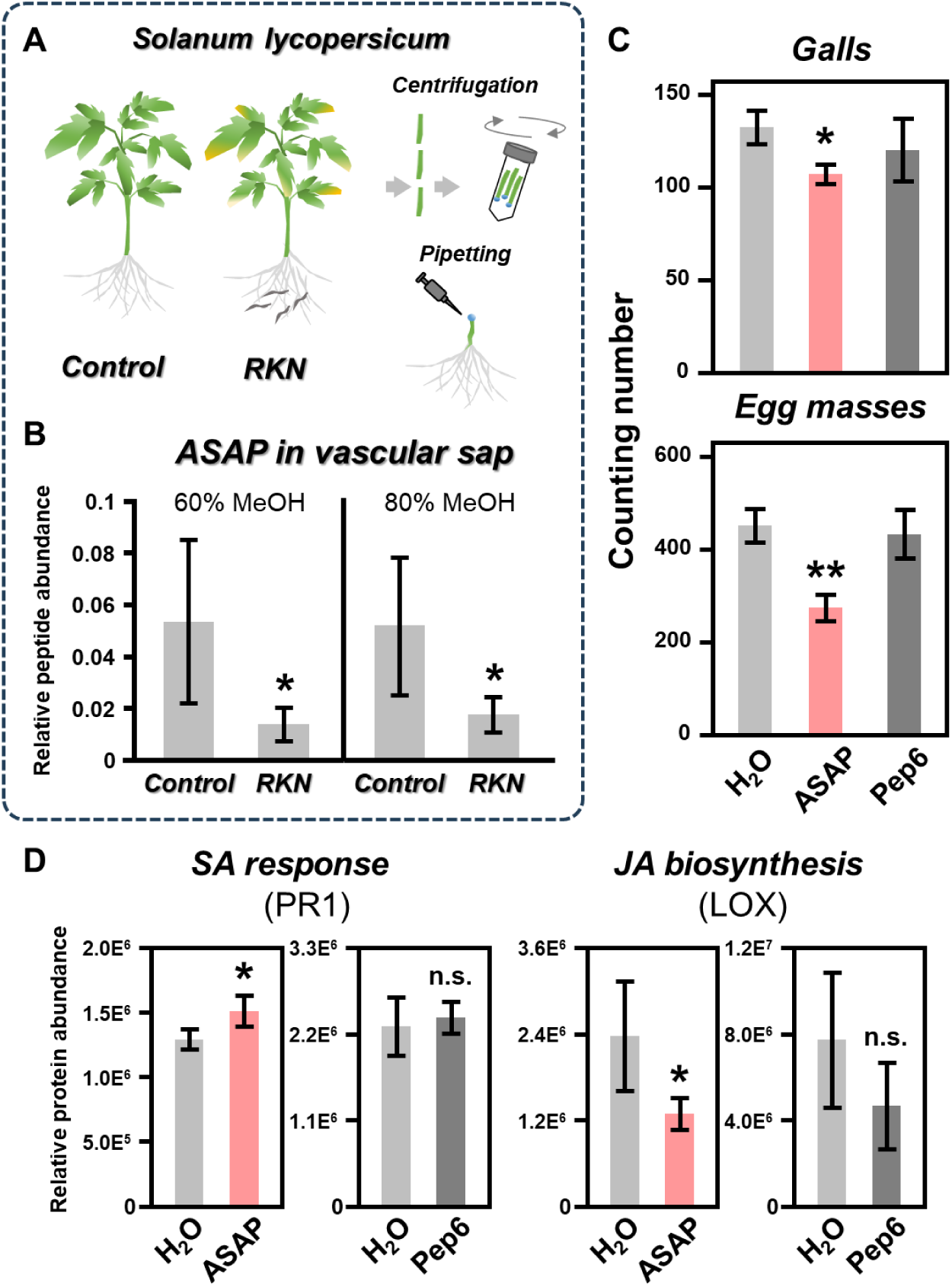
Plant immunity positively regulated by ASAP. (A) The vascular sap collection of the plants with root-knot nematodes (RKN) or without (control) inoculation for ASAP abundance. Vascular sap collection of *Solanum lycopersicum* using both stem centrifugation and pipetting methods. (B) Relative ASAP abundance in the vascular sap of plants with or without RKN inoculation for 7 days were quantified by LC-MS/MS analysis. The extraction of vascular sap peptidome was performed by 60% and 80% methanol (MeOH) elution, respectively. (C) Nematode resistance phenotypes regulated by ASAP treatment. Plants pre-treated with water (control), 1 μM ASAP or 1 μM Pep6 treatment for 24 hours and then inoculated with 500 larvae at J2 stage RKN. The numbers of galls or egg masses were counted in the RKN-inoculated plant roots for 12-day and 45-day post-inoculation, respectively. (D) The protein abundances involved in the SA response and JA biosynthesis regulated by ASAP. PR1, pathogenesis-related protein 1. LOX, lipoxygenase. One and two asterisks represent Student’s *t*-test p < 0.05 and 0.01, respectively. n.s., not significant. Three individual plants were performed as biological replicates for each treatment.

Regarding the conservation of ASAP sequence and its functions across angiosperms, we next explored the conservation of ASAP sequence and its flanking sequences, together as ASAP region, across green plants (Fig. S6A; Table S10). We found that ASAP region are almost identical in angiosperms and gymnosperms (*Pinus taeda* and *Gnetum montanum*), and remained highly conserved in lycophytes (*Selaginella moellendorffii*), moss (*Physcomitrium patens*) and liverwort (*Marchantia polymorpha*) (> 90% sequence identity) (Fig. S6B). In contrast to land plants, ASAP region is much less conserved in algae *Klebsormidium nitens* (21/31 = 68%) and *Chlamydomonas reinhardtii* (16/31 = 52%) (Fig. S6B). In summary, we identified ASAP as a novel peptide family with sequence identical across angiosperms and sequence highly conserved in land plants. In angiosperms, ASAP exerts signaling to regulate lignin biosynthesis, plant growth, and plant immunity through the phosphorylation of a series of MAPKs toward two master regulators.

## Discussion

The conservation of peptide sequences and their signaling pathways has long been considered as a critical issue for understanding how the conserved peptide family regulates essential functions involved in development patterns or defense mechanisms across various species. Peptide families such as CLE, RALF and Pep are reported to be evolutionarily conserved peptides^16, 17, 58, 59^, and exert significant roles in various biological processes. A common method used to identify the conserved peptides across different species typically involves characterizing a peptide in one species through bioassay-guided screening. Subsequently, conserved peptides in other species are often identified solely by *in silico* sequence similarity on the precursor genes. For example, Pep1 was reported in *Arabidopsis thaliana* as an immune-regulatory peptide^60^. Similar sequences of Pep1 were sought in maize and tomato, and synthesized for functional assays without the validation of direct detection on these peptides *in planta*^61, 62^. However, the identification of conserved peptides solely based on the sequence similarity of their precursor genes may introduce significant biases, which could stem from inaccuracies in genome structural annotation. Furthermore, the peptides may not be produced due to the absence of corresponding conserved proteases for cleavage. Thus, the conservation of peptides must be investigated by identifying peptides *in planta* through peptidomic analyses in the species of interest. Previous studies lack comprehensive peptidomic information on the exploration of peptide conservation, which has hindered a systematic investigation of their evolutionary roles in plants. Here, we demonstrated the first successful case on the comprehensive peptidomic analyses in vascular sap throughout six species from the representative evolutionary clades in angiosperms.

Through our peptidomic analyses, ASAP has been identified in six angiosperm species with identical sequences, and conservation analysis based on sequence similarity revealed 100% identity of ASAP in all 114 examined angiosperms (Fig 2B). Compared to all of the known conserved peptide families, ASAP represents the most conserved peptide family to date. Essential housekeeping genes required for maintaining life are also highly conserved, with studies showing approximately 95% identity across 21 species^63^. We thus speculate that the high conservation of ASAP may be essential for maintaining essential functions in plant growth. Given that roots, stems, and leaves are indispensable organs for angiosperm growth, we observed that ASAP can regulate the growth of these organs through phenomic assays. Promoting stem and leaf growth can optimize sunlight exposure for photosynthesis, while root growth facilitates nutrient absorption. Therefore, the regulation of ASAP in roots, stems, and leaves resembles the importance of housekeeping genes in maintaining basic life functions.

Comparing to angiosperms, algae and mosses both lack roots, stems and leaves. We selected four representative species from algae, moss and liverwort (Fig. S6A), and found that ASAP sequences were 100% identical in these species but not in the uni-cellular algae (Fig. S6B). Although multi-cellular algae and early land plants (moss and liverwort) lack roots, stems and leaves, these species possess structures similar to these organs, such as three-dimensional (3D) leafy gametophores or thalli^64–66^. In addition, a previous study reported that the conserved CLE family in moss participates in the transition from 2D precursor tissue to 3D leafy gametophores^67^. Therefore, we hypothesize that the 100% identity of ASAP in land plants and multi-cellular algae could regulate the growth of leaves or leaf-like structures.

Previous studies demonstrated that functionally crucial phosphorylation sites need to be highly conserved to ensure evolutionary functionality. These highly conserved phosphorylation sites are referred to as “phosphorylation hotspots”^68^. This study analyzed phosphoproteomes of 40 eukaryotic species, including humans, mice, nematodes, *Arabidopsis*, and algae, totaling over 530,000 phosphosites, to identify such hotspots^68^. Such analyses revealed that these sites, besides being highly conserved, are often located near catalytic residues or residues involved in protein-protein interactions, mostly in positions crucial for regulatory functions^68^. Mitogen-activated protein kinases (MAPK) could serve as an example for the association between conserved phosphosites and catalytic residues responsible for phosphorylating other proteins. Dual phosphorylation sites on MPK6 forming the TxY motif have been reported to induce structural changes and facilitate substrate recognition^69^. In this study, we observed that ASAP induces phosphorylation of MPK6 also at the TxY motif, and this motif’s sequence is highly conserved across the six species where ASAP was detected (Fig. 3E).

Additionally, the phosphorylation site T927 on the AHA2 homolog in *P. trichocarpa* is highly conserved across the six species (Fig. S2C), and this site has been reported in *Arabidopsis* as T924 in AHA2, which involves the binding of the fungal toxin fusicoccin to AHA2^70^. Since fusicoccin is a plant toxin for the induction of cell wall acidification and leads to plant death upon binding, this conserved phosphosite could be related to ASAP’s regulation of plant immune responses against pathogens.

Several other phosphosites regulated by ASAP on proteins such as SND1, SIRK1, MPK3, and MC4 were also found to be highly conserved (Fig. S2C). While these sites have not been reported before, it is evident that in addition to the conservation of its peptide signal, ASAP plays a conserved functional regulatory role in downstream signaling through these highly conserved regulatory sites.

Plant immunity and growth have always been two crucial aspects in crop research. Initially, it was widely believed that enhancing disease resistance would suppress plant growth. Very few recent studies have shown that proper switching between disease resistance and growth can enhance immunity without affecting growth^71, 72^. The mechanisms involved mostly are related to reversible post-translational modifications (e.g., phosphorylation) for such quick switching. For example, phosphorylated IPA1 was found to induce immunity, and its dephosphorylated form promotes growth^72^. BAK1 is an essential co-receptor for many hormones and peptides, and can regulate growth or immune responses through switching between different phosphorylation sites^71^. Although TFs (IPA1) and co-receptors (BAK1) have shown promising capability in bypassing the immunity and growth trade-off, current known peptides that simultaneously regulate growth and stress responses still face this trade-off dilemma. For example, PSY^73^, PSK^74^, and the RALF family^75, 76^ (e.g. RALF17^77^) are peptides known to inhibit immune responses during growth regulation or suppress growth during defense against pathogens. ASAP, identified in this study, is currently the only peptide found to promote both growth and immune responses.

According to our phosphoproteomic results, ASAP regulates the phosphorylation of TFs related to immune responses (e.g., WRKY33) and growth (e.g., SND1) at different time points post-treatment (Table S5B). Therefore, we speculate that ASAP can modulate the phosphorylation of downstream proteins, activating immune responses when needed and promoting growth when immune responses are not required. This mechanism allows for the optimal balance between immune responses and growth regulation.

## Materials and Methods

### Plant materials and growth condition

Ten to twelve-month-old *P. trichocarpa* and eight-month-old *E. grandis* plants (∼1-1.5 cm diameter of stem) used for vascular sap collection were propagated from the branch cuttings. The rooted branch cuttings were planted into soil and grown in a walk-in chamber for *P. trichocarpa* (16/8 h light/dark cycle at 24-26℃) or in a glasshouse for *E. grandis* at National Taiwan University (Taipei, Taiwan). Twelve-month-old *C. kanehirae* used for sap collection were grown from the ten-year-old tree stumps at Xinxian nursery, Taiwan forestry research institute, Council of Agriculture (Taipei, Taiwan). Five to six-week-old *S. lycopersicum* (cv. CL5915) used for sap collection were grown from the seeds. The seeds were immersed in ddH_2_O for 48 hours at 4℃, and then transferred into soil for growing in a walk-in chamber (12/12 h light/dark cycle at 24-26℃) at National Cheng Kung University (Tainan, Taiwan). Two to three-week-old *Zea mays* (cv. White Pearl) and two to three-week-old *Glycine max* (L. cv. Kaohsiung selection no.10) used for sap collection were grown from the seeds. The seeds were sown into soil and grown in a phytotron (16/8 h at 25/23℃ of light/dark cycle) at CH Biotech R&D Co., LTD (Nantou, Taiwan).

Four-month-old *P. trichocarpa* plants were grown and used for ASAP-induced monolignol metabolite detection and time-course phosphoproteomic analysis. Two-month-old *E. grandis* plants were grown for monolignol metabolite detection. Nine-month-old *P. trichocarpa* were used for detection of ASAP migration in vascular sap and stem-differentiating xylem. These plants used for ASAP treatment were grown from branch cuttings and maintained in the same glasshouse mentioned above in National Taiwan University.

Eight-day-old *Arabidopsis thaliana* and three-week-old *S. lycopersicum* seedlings used for lignin staining experiment were grown on agar plate containing 1/2 Murashige and Skoog medium (Duchefa Biochemie) with 1% sucrose and 0.8% agar. The seeds of *A. thaliana* (Col-0) and *S. lycopersicum* (cv. CL5915) were surface-sterilized with 1.5% bleach before use. These seeds were grown on agar plates at 4℃ for 2 days, and then the agar plates were transferred to a growth chamber (16/8 h light/dark cycle at 22℃).

Two-month-old *P. trichocarpa* and two-week-old *Z. mays* grown in the glasshouse at National Taiwan University and six-week-old *S. lycopersicum* grown in a walk-in chamber (12/12 h light/dark cycle at 24-26℃) at National Cheng Kung University were transferred to the phenotyping greenhouse at Taiwan Agricultural Research Institute for high-throughput phenomic analyses.

Fourteen-day-old of *S. lycopersicum* (cv. CL5915) seedlings used for root biomass observation were grown in growth chamber (12/12 h light/dark cycle at 25℃) at Taiwan Agricultural Research Institute. The seeds were immersed in ddH_2_O for 7 days until rooting, and then transferred into Kimura culture medium^78^ for growing 7 days.

Four to five-week-old *S. lycopersicum* (cv. CL5915) used for inoculation with root-knot nematodes and examination of ASAP-induced immunity were grown from seeds and maintained in the same walk-in chamber mentioned above at National Cheng Kung University.

### Vascular sap collection

For peptidomic profiling, the vascular sap samples of *Z. mays*, *S. lycopersicum*, *P. trichocarpa*, *G. max* and *E. grandis* were collected by pipetting method. Stems of these plants were cut from about 2 cm above soil, and then the sap accumulated in the top of each stump was collected by pipetting for around 30 minutes. The vascular sap sample of *C. kanehirae* was collected by centrifugation method. The branches (1 cm diameter; about 8-9 cm length) were cut and placed into 50-ml tubes followed by centrifugation at 3,000 ×g at room temperature to collect the sap. For targeted detection of endogenous ASAP or isotope-labeled ASAP, the vascular sap samples of *P. trichocarpa* or *S. lycopersicum* were collected from the stem segments by centrifugation at 3,000 ×g at room temperature. To confirm that the 3,000 ×g centrifugation would not damage the cells, the *P. trichocarpa* protoplasts were used for centrifugation by the same condition (Fig S7). We found that the number of protoplasts was not significantly changed after 3,000 ×g centrifugation twice. The detection of ASAP from vascular sap using two different collection methods was showed in Fig. S1. The vascular sap samples were collected from three individual plants for biological replicates to detect the endogenous ASAP and isotope-labeled ASAP. All sap samples were frozen in liquid nitrogen and then stored at −80℃ before peptide purification.

### Vascular sap and xylem peptide purification

Peptides were extracted and purified from each sap sample using the Sep-Pak tC18 columns (Part No. WAT036795, Waters, 1 g, 6 cc). The columns were conditioned with 10-ml 100% methanol and then equilibrated with 15-ml 0.1% trifluoroacetic acid (TFA). The collected sap was loaded into the column and then was washed by 10-ml 0.1% TFA. Total 15-ml 60% methanol and 15-ml 80% methanol were sequentially used to elute the retained peptides on the columns. The 60% and 80% methanol-eluted samples were dried by a vacuum centrifugation concentrator (miVac Duo Concentrator; Genevac), then the dried samples were dissolved in 100 to 400 μl 0.1% TFA. The supernatant was transferred to a new tube and the undissolving pellet was removed after 13,000 ×g centrifugation for 10 min at 4℃. The supernatant was desalted by the ZIPTIP® C18 pipette tips (Millipore) and eluted by 50% acetonitrile (ACN) in 0.1% TFA. The desalted samples were dried by a vacuum centrifugation concentrator, and then dissolved in 0.1% formic acid (FA) and centrifuged at 13,000 ×g for 10 min at 4℃. The supernatant was transferred into a sample vial, and analyzed by liquid chromatography tandem mass spectrometer (LC-MS/MS).

To detect the isotope-labeled ASAP in living xylem, the xylem sample was collected into liquid nitrogen by scraping the surface of the debarked stems. Before peptide extraction, xylem sample was ground into fine powder under liquid nitrogen. Total 2.5 g powder of each sample was resuspended in 30-ml 1% TFA with 250 μl protease inhibitor cocktail (Roche) and then centrifuged at 12,000 ×g for 20 min at 4℃. The supernatant was transferred to the new 50-ml tube and the pH value was adjusted to 4.5 by NaOH. The debris was removed by 12,000 ×g centrifugation for 20 min at 4℃. The supernatant was transferred again to a new 50-ml tube and the pH value was adjusted to 2.5 by HCl. Peptides were extracted and purified from each sample using the Sep-Pak C18 columns and ZIPTIP® C18 pipette tips following the same processes as the sap samples described above.

### LC-MS/MS-based sap peptidomic analyses

For sap peptidomic profiling, the 60% and 80% methanol-eluted sap samples were analyzed by a Dionex 3000 UPLC system coupled with a Q Exactive Hybrid Quadrupole-Orbitrap mass spectrometer (Thermo Fisher Scientific). Peptides were separated on a 25-cm PepMap C18 column packed with 2-μm particles (Thermo Fisher Scientific) using the mobile buffer consisted of 0.1% FA in ultra-pure water with an eluting buffer of 0.1% FA in 100% ACN with a 60 min linear gradient of 5 to 25% ACN/0.1% FA at a flow rate of 300 nl/min.

For *S. lycopersicum*, the 60% and 80% methanol-eluted sap samples were analyzed by a Dionex 3000 UPLC system coupled with a Q Exactive Plus Hybrid Quadrupole-Orbitrap mass spectrometer (Thermo Fisher Scientific). Peptides were separated on a 25-cm PepMap C18 column packed with 2-μm particles (Thermo Fisher Scientific) using the mobile buffer consisted of 0.1% FA in ultra-pure water with an eluting buffer of 0.1% FA in 100% ACN with a 90 min linear gradient of 11 to 37% ACN/0.1% FA at a flow rate of 300 nl/min.

The mass spectrometer was operated in the data-dependent acquisition (DDA) mode, in which a full scan (from m/z 350-1600 with the resolution of 70,000 at m/z 200) was followed by higher-energy collisional dissociation fragmentation using the 20 most intense ions with normalized collision energy of 27 and the resolution of 17,500 at m/z 200. The automatic gain control (as AGC) value of MS was set as 1e5 with the max injection time as 120 ms. The isolation window was set as 2.0 while the dynamic exclusion duration as 20 s. The raw data of sap peptidome was processed into a format suitable for MS/MS ion search against target-decoy database.

To increase the detection sensitivity of endogenous ASAP and isotope-labeled ASAP using the same instrument, the parallel reaction monitoring method was applied to the targeted m/z 549.7834 and m/z 552.7874 for ASAP and isotope-labeled ASAP, respectively.

For *Z. mays* and *G. max*, the 60% and 80% methanol-eluted sap samples were analyzed by an Easy nLC 1200 coupled with Orbitrap Fusion Lumos mass spectrometer (Thermo Fisher Scientific). Peptides were separated by an Acclaim PepMap 100 C18 trap column (75 µm x 2.0 cm, 3 µm, 100 Å, Thermo Fisher Scientific) and an Acclaim PepMap RSLC C18 nano LC column (75 µm x 25 cm, 2 µm, 100 Å) using the mobile buffer consisted of 0.1% FA in ultra-pure water with an eluting buffer of 0.1% FA in 100% ACN with 5 to 25% ACN/0.1% FA at a flow rate of 300 nl/min. The 90 min linear gradient was used. For DDA mode, a full scan (m/z 350-1600 with the resolution of 120,000 at m/z 200) was followed by MS/MS fragmentation in a 3 sec cycle time. The maximum ion injection time was set 50 ms. The MS/MS acquisitions were performed using 1.4 Da isolation window with 50,000 AGC value, 35% normalized collision energy, maximum IT 120 ms, and 15,000 resolving power.

### Protein extraction, digestion and phosphopeptides enrichment

The debarked stems of *P. trichocarpa* plants with or without ASAP treatment for different time points (0.5, 1, 2 and 4 hours) were used to extract total proteins using a previous published method^79^. Protein pellet of each sample was dissolved in 5% SDS/50 mM Tris-HCl (pH 8.0), and then the concentrations of dissolved proteins were measured using BCA protein assay (Thermo Fisher Scientific). Before trypsin digestion, the disulfide bonds of proteins were reduced with 10 mM tris (2-carboxyethyl) phosphine and alkylated with 40 mM chloroacetamide at 45 ℃ for 10 min. Each protein sample was digested into peptides by trypsin in a S-trap micro column to remove the incompletely digested proteins and trypsin. The tryptic peptides were acidified by TFA and desalted by SDB-XC StageTips (catalog no. 2340; 3M). The desalted samples were dried by a vacuum centrifugation concentrator. The dried samples were dissolved in 0.1% formic acid and centrifuged at 13,000 ×g for 10 min at 4℃. The supernatant was transferred into a sample vial, and analyzed by for LC-MS/MS analysis.

To enrich the phosphopeptides from total tryptic peptides, the immobilized metal ion affinity chromatography (IMAC) StageTip was made and performed for phosphopeptide enrichment following by a previous reported method^80, 81^. The tip-end was capped with a 20-μm polypropylene frits disk (Aglient) and then packed with Ni-NTA silica resin (QIAGEN, Hilden, Germany) by centrifugation at 200 ×g for 1 min. The Ni^2+^ ions of resin within a tip were removed by adding 100 μl of 100 mM EDTA and then coupled with Fe^3+^ by adding 100 μl of 100 mM FeCl_3_. The Fe^3+^-NTA IMAC StageTips were made and equilibrated with 6% acetic acid at pH 3.0 before sample loading. The tryptic peptides were dissolved in 6% acetic acid at pH 3.0 and loaded into the IMAC StageTip, and then washed the non-phosphopeptides using 4% acetic acid in 25% ACN and equilibrated with 6% acetic acid. After washing, the IMAC-bound phosphopeptides were eluted by 150 μl of 200 mM NH_4_H_2_PO_4_. The eluted phosphopeptides were desalted using SDB-XC StageTips (catalog no. 2340; 3M). The dried samples were dissolved in 0.1% formic acid and centrifuged at 13,000 ×g for 10 min at 4℃. The supernatant was transferred into a sample vial, and analyzed by for LC-MS/MS analysis.

### LC-MS/MS-based phosphoproteomic analyses

The enriched phosphopeptides were analyzed by an Easy nLC 1200 coupled with Orbitrap Fusion Lumos mass spectrometer (Thermo Fisher Scientific). Peptides were separated by an Acclaim PepMap 100 C18 trap column (75 µm x 2.0 cm, 3 µm, 100 Å, Thermo Fisher Scientific) and an Acclaim PepMap RSLC C18 nano LC column (75 µm x 25 cm, 2 µm, 100 Å) using the mobile buffer consisted of 0.1% FA in ultra-pure water with an eluting buffer of 0.1% FA in 100% ACN with 5 to 25% ACN/0.1% FA at a flow rate of 300 nl/min. A 90 min linear gradient and DDA method was used. The DDA method contains a full MS scan (m/z 350-1600 with the resolution of 120,000 at m/z 200) following by MS/MS fragmentation in a 3 sec cycle time. The maximum ion injection time was set 50 ms. The MS/MS acquisitions were performed using Da isolation window with 50,000 AGC value, 35% normalized collision energy, maximum IT 120 ms, and 15,000 resolving power.

### Target-decoy database for peptide-spectrum matching

Target-decoy database was used for MS/MS ion searching and false discovery rate (FDR) evaluation to identify endogenous peptides^82^, which was generated by combining the protein sequences with the randomized protein sequences (decoy) for each species using Trans Proteomics Pipeline (TPP) version 5.1^83^. All protein fasta files of six species (*Zea mays* Zm-B73-REFERENCE-NAM-5.0.55, *Cinnamomum kanehirae* v3, *Solanum lycopersicum* ITAG5.0, *Eucalyptus grandis* v2.0, *Glycine max* Lee v1.1 and *Populus trichocarpa* v4.1) were downloaded from Phytozome^84^.

### MS data processing and database searching

For peptidomic profiling, the MS raw data obtained from LC-MS/MS was first converted into mzXML format using MSConvert^85, 86^, and then processed by UniQua with optimized parameters (Smoothing Width: 5, Centroiding, Deisotoping, Noise Removal, Intensity Cutoff: 500)^87^. The UniQua processed spectra were converted into Mascot generic format (.mgf) by MSConvert^86^. The processed mgf files were searched against a concatenated target-decoy database for corresponding species (described above) using Mascot search engine (version 2.3; Matrix Science, London, UK) without specifying enzyme cleavage rules. The mass tolerances used in the MS/MS ion search for peptide precursors and fragments were 10 ppm and 0.05 Da, respectively. The oxidation of methionine and the sulfation of tyrosine were considered as the variable modifications. Three independent scoring methods, Mascot, Delta Score (DS), and Contribution Score (CS)^29^ were used to evalute the peptide identification based on FDR. The Mascot search result files (.dat) were used to extract the Mascot scores and to re-calculate the scores for DS and CS based on Mascot scores using the CScore program^29^. The identified peptides with FDR below 0.05 from at least one of the three scoring methods were regarded as positive hits.

For phosphoproteomic analysis, the MS raw data were searched against protein database using Sequest, Mascot and Byonic search engines in Proteome Discoverer (version 2.5). Peptide precursor mass tolerance was set at 10 ppm, and MS/MS tolerance was set at 20 ppm. A static modification of carbamidomethylation on cysteine residue (+57.0214 Da) and variable modifications of oxidation on methionine residue (+15.9949 Da) and phosphorylation on serine, threonine, and tyrosine residues (+79.9663 Da) were set for search. Searching was performed with full tryptic digestion and allow a maximum of two missed cleavages on the peptides. The criteria of FDR were set as 0.01 for both peptide and protein identification. Label free quantification was performed using the nodes of Minora feature detector and precursor ions quantifier in Proteome Discoverer.

A phosphopeptide was considered as significant change in its phosphorylation state through *p*-value < 0.05 based on one-sided Student’s *t*-test and with expression fold change greater than (up-regulated upon ASAP treatment) or lower than 0.67 (down-regulated upon ASAP treatment). For the *t*-tests, variances of control group and ASAP treatment group were assumed to be different if the *p*-values of the F-tests were smaller than 0.05. Fold change was calculated as the average expression level of the three biological replicates of ASAP treatment group divided by the average expression level of the three biological replicates of control group.

### Peptidomic comparative conservation analysis

Total peptides identified in one species were used as target sequences to search against the protein sequences of each other five species using BLASTP^88^ (e-value < 100 and default settings). One peptide was considered as conserved if the full length of this peptide could be completely aligned to a protein sequence from the other species and the sequence identity was equal to or more than 65%. The protein fasta files were downloaded from Phytozome as described above^84^.

### Position weight matrix (PWM) analysis

The potential ASAP candidates across angiosperms were examined using the 114 species selected from 50 families in distinct clades of angiosperms. All protein fasta files were downloaded from Phytozome or the National Center for Biotechnology Information (NCBI) (Table S4). The sequence of ASAP and its 10 flanking residues on both N-terminal and C-terminal sites were extracted from all 114 species and applied for PWM analysis. Each species was identified with at least one ASAP sequence. PWM of ASAP and its flanking sequences was generated by the default settings of WebLogo (version 2.8.2)^89^.

To examine the conservation of phosphorylation sites (later as phosphosites), the identified phosphosite with its 5 flanking residues on both sites (here designed as phosphosite region) of SIRK1, AHA2, MPK3, MPK6, MC4 and SND1 in *P. trichocarpa* were applied for PWM analysis. Sequences of phosphosite regions were queried against the protein sequences of each remaining five species using BLASTP with default settings^88^. For each phosphosite region, the best hit with the highest identity under full-length alignment was selected in each species. Each phosphosite region, along with their BLASTP best hit sequences, were aligned using ClustalW within MEGA-X under default settings^90, 91^. Aligned results were used as input to generate PWM using WebLogo (version 2.8.2)^89^.

### Synthetic peptide treatment

Synthesized ASAP (XXXXXXXXXXX, averaged MW: 1098.205), the powder with ∼95% purity was purchased from GeneScript. The synthesized isotope-labeled ASAP (XXXXXX[X(^13^C_5_, ^15^N_1_)]XXXX, averaged MW: 1104.224) powder with ∼95% purity was purchased from GeneScript. The other synthesized peptides, flg22 (QRLSTGSRINSAKDDAAGLQIA, averaged MW: 2272.483) and Pep6 (ATDRRGRPPSRPKVGSGPPPQNN, averaged MW: 2441.674), the powder with ∼95% purity was purchased from Mission Biotech Inc.

A stock solution of ASAP (1 mg/ml) was prepared by dissolving 1 mg ASAP powder in 1 ml ddH2O and stored at −80℃. The purity and molecular weight of ASAP in the stock solution were further examined by the ACQUITY UPLC system coupled with Synapt G2 HDMS (Waters) (Fig. S8). The different concentrations (0.1, 0.5 and 1.0 µM) of ASAP working solutions were freshly prepared by dilution from the stock solution. For phosphoproteomic analysis, 50 ml of 1.0 µM ASAP solution was irrigated to the plants for collecting samples at 0.5, 1, 2 and 4 hours. The same volume of ddH_2_O was irrigated to the plants as the control group for each time point. For ASAP-treated phenotypic changes and monolignol biosynthesis, 0.1, 0.5 and 1.0 µM of ASAP solution were irrigated to the plants three times per week (50 ml per time and 100 ml per time in the last three times) for around one month. The same volume of ddH_2_O was irrigated to the plants as the control group with the same frequency.

The stock and working solutions of isotope-labeled ASAP were prepared as described above. For the detection of exogenous isotope-labeled ASAP in vascular sap and living xylem of plants, 100 ml of 1.0 µM isotope-labeled ASAP solution was irrigated to the plants for 2 days to collect the samples. For lignin staining, 1.0 or 2.0 µM ASAP or 1.0 µM flg22 solutions in 1/2 Murashige and Skoog medium were treated to seedlings for 24 hours. For induction of plant immunity, 1.0 µM ASAP or 1.0 µM Pep6 were irrigated to the plants.

### Metabolite extraction

The stems of ASAP- or ddH_2_O-treated plants used for compound extraction were ground into fine powder in liquid nitrogen. The 0.3 g frozen powder of each sample was mixed with 0.9 ml methanol-chloroform (2:1) solution pre-cooled at −20℃, and then rotated gently for 10 min at 4℃. After centrifugation at 13,000 ×g for 10 min at 4℃, the supernatant was transferred to new tubes, and 0.1 ml ddH_2_O was added. The ddH_2_O-methanol-chloroform mixture was rotated for 10 min at 4℃ and then centrifuged at 13,000 ×g for 10 min at 4℃ to separate two liquid layers. The upper (hydrophilic) layer and the lower (hydrophobic) layer were collected separately into new tubes, and then dried by a vacuum centrifugation concentrator (miVac Duo Concentrator; Genevac). The pellets from upper and lower layers were dissolved with 100 μl 4℃ pre-cooled methanol-ddH2O (1:1) and 300 μl −20℃ pre-cooled chloroform-methanol (2:1) solution, respectively. Both re-dissolved upper- and lower-layer solutions were centrifuged at 13,000 ×g for 10 min at 4℃, and the corresponding supernatants were transferred into sample vials for the quantification of compounds in monolignol biosynthesis pathway.

### Standard compounds of monolignol biosynthesis pathway

Ten standards were used to develop the targeted quantitative method for the compounds in monolignol biosynthesis pathway. Nine standards were purchased, including caffeic acid (98%, AK Scientific), ferulic acid (99%, Acros Organics), 4-coumaryl alcohol (98%, Toronto Research Chemicals), coniferaldehyde (99.94%, MedChemExpress), coniferyl alcohol (97.5%, Supelco Merck), 5-hydroxyferulic acid (95%, Sigma-Aldrich Merck), sinapic acid (98%, AK Scientific), sinapaldehyde (99.96%, MedChemExpress) and sinapyl alcohol (95%, Biosynth). One remaining standard, 5-hydroxyconiferaldehyde (5H-Ald), was synthesized in this study (the details was shown below).

The structural characteristics of synthetic 5H-Ald were examined by nuclear magnetic resonance (NMR), high-resolution mass spectrometry (HRMS) and high-performance liquid chromatography (HPLC). Proton (^1^H) and carbon (^13^C) NMR spectra were recorded on a Bruker Avance III 400 (400 MHz for ^1^H; 100 MHz for ^13^C) using MeOD-*d_4_* as solvent. Chemical shifts were reported in δ (parts per million, ppm) using the residual solvent (MeOD-*d_4_*, 3.31 ppm) as reference standard. The coupling constants were given in hertz (*J*/Hz). High-resolution mass spectrum was recorded on a Q Exactive Plus Hybrid Quadrupole-Orbitrap mass spectrometer (Thermo Fisher Scientific). The purity of 5-hydroxyconiferaldehyde was more than 95%, which was validated by the Agilent Technologies HPLC system (1260 Infinity II) using C-18 column (Kinetex® 2.6µm EVO C18 100 Å, 100 × 4.6 mm). The spectra of NMR and HRMS and the chromatogram of HPLC were shown in Fig. S9A to D.

### Quantitation of compounds in monolignol biosynthesis pathway by LC-MS/MS

A mixture of standards (50 ng/µl of each standard in methanol) was analyzed using the Vanquish UHPLC system coupled with a Dual-Pressure Ion Trap Mass Spectrometer (Velos Pro, Thermo Fisher Scientific). Each compound was separated by an HSS T3 column (Waters ACQUITY HSS T3 100Å, 1.8 µm, 100 x 2.1 mm) at 40℃ using the mobile buffer consisted of 2% ACN/0.1% FA (Buffer A) with an eluting buffer of 100% ACN/0.1% FA (Buffer B) with a 11 min gradient of 0.5-30% Buffer B at 0 to 6 min, 30 to 50% Buffer B at 6 to 7 min, 50 to 99.5% Buffer B at 7 to 7.5 min, 99.5% Buffer B at 7.5 to 9.5 min, 99.5 to 0.1% Buffer B at 9.5-10 min and then equilibrated by 0.1% Buffer B at 10-11 min. The mobile phase without FA was specifically applied for the detection of 4-coumaryl alcohol and coniferyl alcohol.

The mass spectrometer was operated in the negative ion mode with one full MS scan (m/z 100 to 350), then switched to seven selected reaction monitoring (SRM) scans for each compound: m/z of 179.03 to 135 for caffeic acid, 193.05 to 149 for ferulic acid, 177.05 to 162 for coniferaldehyde, 209.05 to 194 for 5-hydroxyferulic acid, 223.06 to 208 for sinapic acid, 207.07 to 192 for sinapaldehyde, 193.05 to 178 for 5-hydroxyconiferaldehyde. The SRM scans were set to m/z of 149.06 to 131 for 4-coumaryl alcohol, 179.07 to 161 for coniferyl alcohol and 193.09 to 161 for sinapyl alcohol using the mobile phase without FA. The relative abundance of each compound detected in hydrophilic or hydrophobic layer was obtained by peak area calculation of the selected product ions detected in SRM scans.

### Lignin staining

Seedlings were transferred from agar plates to the 48-well plate (*A. thaliana*) or 24-well plate (*S. lycopersicum*) as one seedling in one well, and each well contained 0.8-ml 1/2 Murashige and Skoog medium. After peptide or ddH_2_O treatment for 24 hours, seedlings were incubated with 40% HCl for 3 to 5 min, and then moved onto a glass slide, then immersed using 5% phloroglucinol in 95% ethanol for around 15 seconds. The lignified tissues were showed as the pink-red coloration and observed using a dissecting microscopy.

### Phenomic analyses

To observe the changes of plant growth, the high-throughput phenomic analyses were performed for recording and calculating the digital biomass of each plant during water (control) or different concentrations of ASAP treatment. The digital biomass was calculated by the multiplication of plant height by leaf area, and these parameters were recorded by a high-throughput multispectral 3D scanner, PlantEye F500 (Phenospex) in the phenotyping greenhouse at Taiwan Agricultural Research Institute, Ministry of Agriculture. These high-throughput phenotypic datasets were further processed by the smoothing and extraction of traits (SET)-based analysis following a previously published method^92^.

To examine the changes in digital biomass under the effect of ASAP treatment, we subtracted the biomass value of the first day of ASAP treatment (day 0) from the value of the following days for each plant. As shown in Fig. S10, the increased biomass of a plant 2 days after 0.5 μM ASAP treatment was calculated by subtracting the day 0 biomass from the day 2 biomass after 0.5 μM ASAP treatment, shown as [Day 2 – Day 0 biomass]_0.5μM_. The increased biomass of the control group was also calculated by subtracting the day 0 biomass from day 2 biomass, shown as [Day 2 – Day 0 biomass]_control_. The fold change of the biomass change after 0.5 μM ASAP treatment for 2 days was calculated by [Day 2 – Day 0 biomass]_0.5μM_/[Day 2 – Day 0 biomass]_control_. The averaged value of the biological replicates was applied for the calculation of fold change using GateMultiplex^93^.

At each time point, outliers were identified from each treatment group and removed independently. An outlier was defined as a value smaller than the first quartile (Q1) minus 1.5 times the interquartile range (IQR), or greater than the third quartile (Q3) plus 1.5 times the IQR^94^.

The biomass changes of the plant roots with ASAP treatment were observed by a high-resolution image scanner (Perfection V850 Pro, EPSON). Total root length, taproot length, lateral root length, and root volume were recorded and analyzed by using RhizoVision Explorer software (v2.0.3)^95^. Outliers were removed using the same criteria as described in the last paragraph.

### Plant resistance assay for root-knot nematodes

The cultivation of root-knot nematodes *Meloidogyne incognita* was performed in four to five-week-old *S. lycopersicum* (cv. CL5915) plants. After 45 days of inoculation, egg masses were harvested from the roots and then hatched in sterilized ddH_2_O in a glass dish under sterile conditions. The resulting J2 larvae of *M. incognita* were counted using a dissecting microscopy. After the larvae collection, each plant with 24 hours of peptide treatment was inoculated with a total 500 larvae of *M. incognita*. The solution with larvae was added into the soil for the inoculation on the plant roots. The number of galls and egg masses were observed after 12 days and 45 days after inoculation.

### Sequence alignment of ASAP and its flanking sequence across green plants

Peptide sequence of ASAP and its 10 flanking residues on both sides were extracted from the six species (*Z. mays*, *C. kanehirae*, *S. lycopersicum*, *E. grandis*, *G. max* and *P. trichocarpa*), and aligned with the same length of sequence region to other species across green plants, including *C. reinhardtii*, *K. nitens*, *M. polymorpha*, *P. patens*, *S. moellendorffii*, *P. taeda* and *G. montanum*. For each species, the BLASTP hit of the six-species consensus ASAP and its flanking sequence with the smallest e-value were displayed in Fig. S6 (Table S10).

The additional protein fasta files were downloaded from Phytozome (*Chlamydomonas reinhardtii* v5.6, *Marchantia polymorpha* v3.1, *Physcomitrium patens* v3.3 and *Selaginella moellendorffii* v1.0)^84^, NCBI (*Klebsormidium nitens*, GenBank accession: GCA_000708835.1), and TreeGenes^96^ (*Pinus taeda* v2.01 and *Gnetum montanum* v1.0).

## Acknowledgments

We thank Ni-Chiao Tsai for assisting the vascular sap collection of *E. grandis*, Wei-Hung Chang for UniQua software operation, Dr. Shu-Ming Tsao from ez-Omics Co., Ltd. for target-decoy FDR calculation, and thank Prof. Pei-Chen Chen for providing root-knot nematodes, Prof. Lay-Sun Ma, Prof. Chia-Lin Chung and Zong-Chi Wu for assistance of pathogen infection test. The mass spectrometry analyses were supported by the MS-based Multi-Omics Platform, University Center of Bioscience and Biotechnology, National Cheng Kung University. The targeted metabolite analyses were supported by the Metabolomics Core Facility at the Agricultural Biotechnology Research Center of Academia Sinica.

## Author Contributions

Y.L.C., Y.J.L. supervised the project and designed the experiments. Y.L.C., P.C.L., Y.F.H., J.H.Y. performed the vascular sap peptide experiments. Y.L.C., P.C.L., C.W.C., C.C.H. performed phosphoproteomic experiments. Y.L.C., P.C.L. performed metabolite experiments. Y.L.C., Y.F.H., C.W.C., C.C.H. performed the LC-MS/MS analysis. Y.L.C., C.H.C. performed MS data processing and analysis. Y.F.H., K.H.H., I.F.W. performed lignin staining experiments. K.X. provided the *Z. mays* and *G. max* samples, C.C.W. provided and collected the *C. kanehirae* samples. J.C.S. performed the monolignol compound synthesis and purity test. Y.K.T., C.B.L., D.G.L. performed phenomic analyses, C.Y.K., Y.F.H. performed root-knot nematode experiments. Y.L.C., Y.J.L., C.H.C., P.C.L. constructed all figures. Y.J.L., Y.L.C., C.H.C., P.C.L., I.J.T. wrote the manuscript.

## Data Availability

All peptidomic and phosphoproteomic data were deposited to the ProteomeXchange Consortium via the PRIDE partner repository with the dataset identifier PXD032088^97^. All other data needed to evaluate the conclusions in the paper are present in the paper and/or the Supplementary Materials.

## Supplementary Figures

**Fig. S1.**
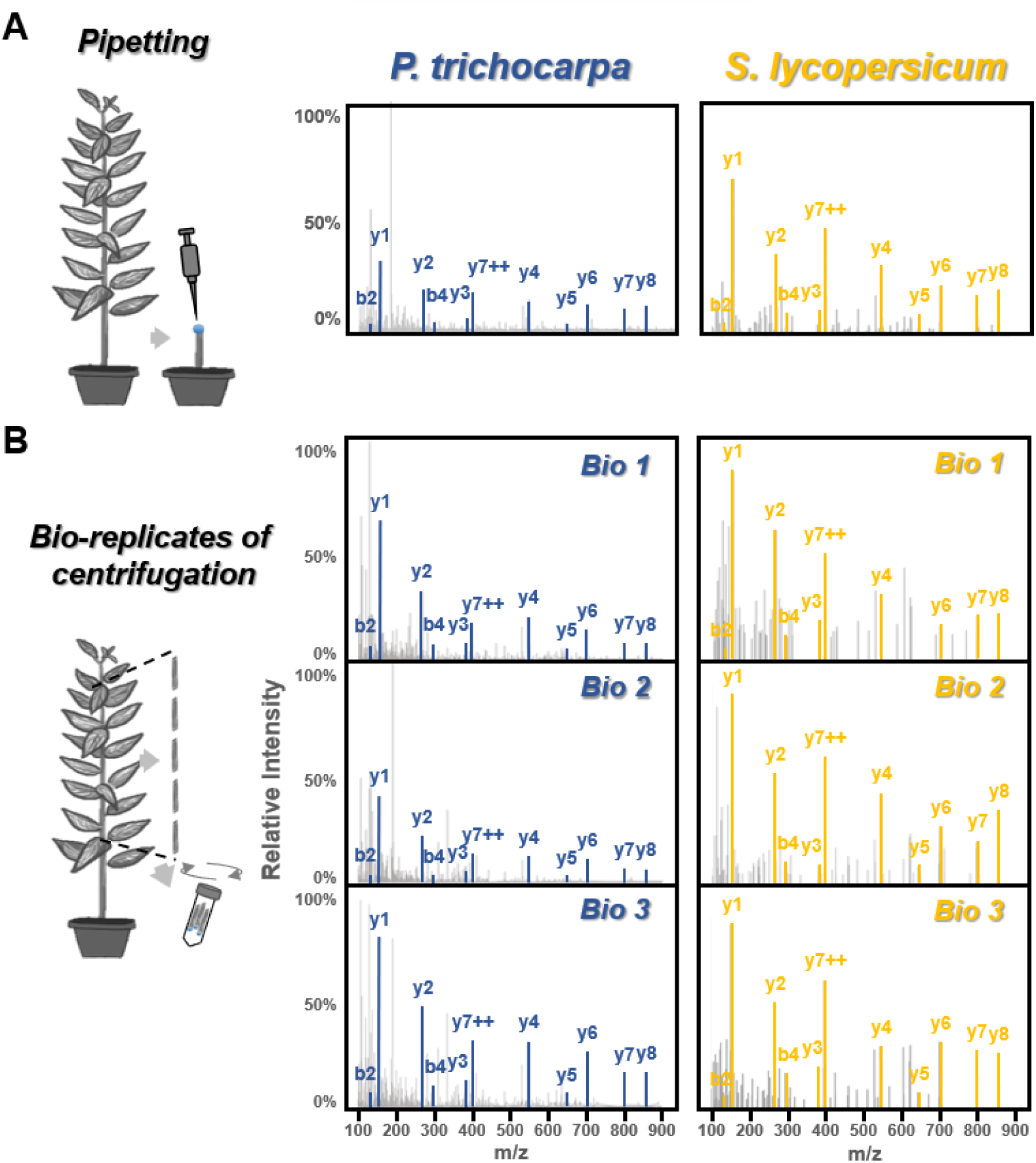
Detection of endogenous ASAP in vascular sap collected by centrifugation or pipetting. For different sap collection methods, the detection of ASAP in the vascular sap of *Populus trichocarpa* and *Solanum lycopersicum* were analyzed by LC-MS/MS. (A) Sap collection by pipetting for ASAP identification. (B) Validation of ASAP using centrifugation method for sap collection for three biological replicates (labeled as Bio 1, 2 and 3). The b and y fragment ions of ASAP generated by MS/MS fragmentation were labeled in all spectra. m/z, mass-to-charge ratio.

**Fig. S2.**
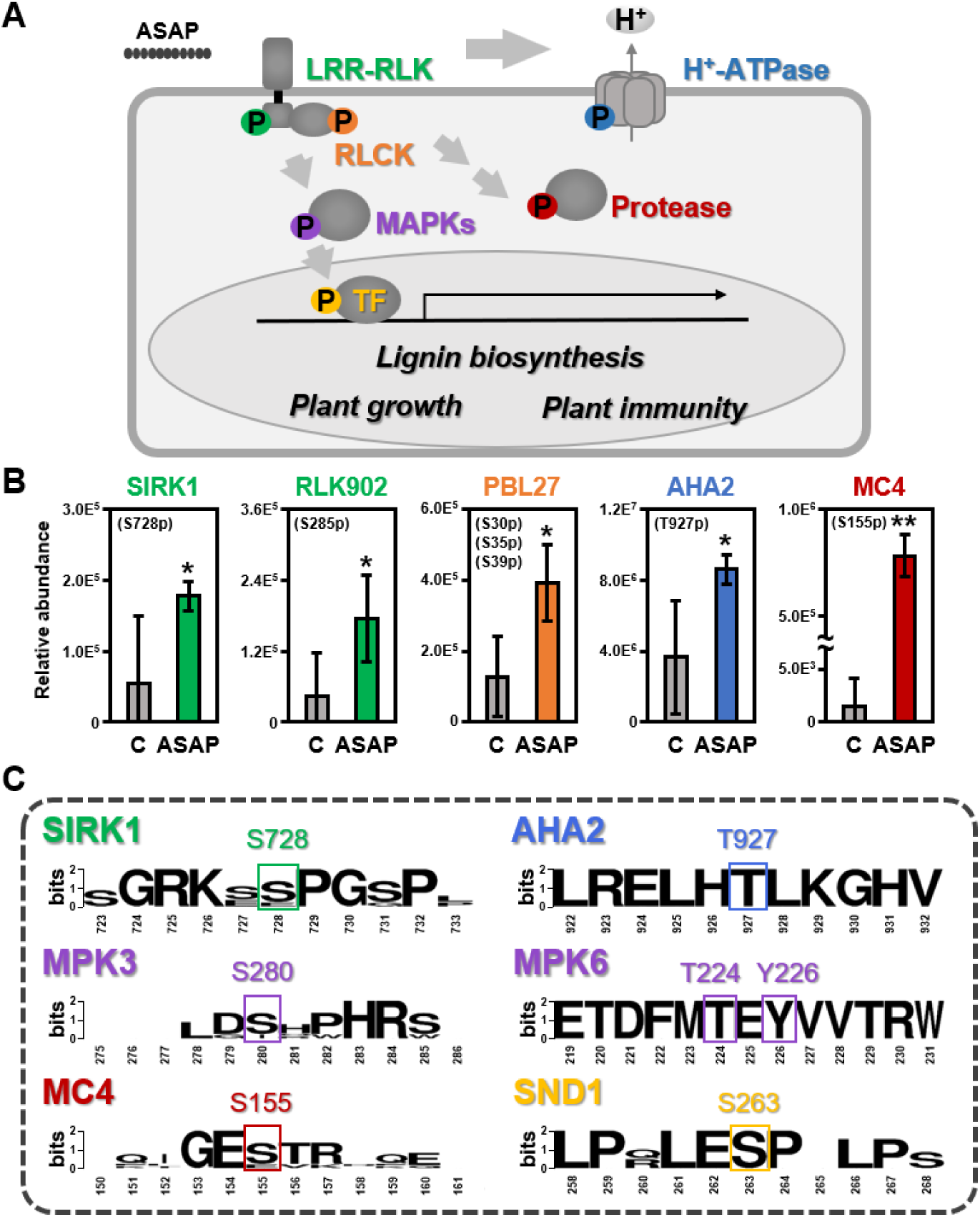
Conserved phosphosites on known functional proteins regulated by ASAP. (A) Proposed model of ASAP signaling based on phosphorylation cascades. LRR-RLKs, leucine-rich repeated receptor-like kinases. RLCK, receptor-like cytoplasmic kinase. MAPKs, mitogen-activated protein kinases; TF, transcription factors. (B) The phosphopeptide levels of selected proteins regulated by ASAP. The debarked stems of *P. trichocarpa* treated by ddH2O (labeled as C) or ASAP were used for phosphoproteomic analyses. (C) Position weight matrix of phosphosite regions on different proteins extracted from 6 species, including *Zea mays*, *Cinnamomum kanehirae*, *Solanum lycopersicum*, *Eucalyptus grandis*, *Glycine max* and *Populus trichocarpa*. SIRK1, sucrose-induced receptor kinase 1. RLK902, receptor-like kinase 902. PBL27, PBS1-like 27. AHA2, AUTOINHIBITED H^+^-ATPase 2. MC4, metacaspase 4. MAPKKK3, MAPK kinase kinase 3. MKK4, MAPK kinase 4. MPK3, mitogen-activated protein kinase 3. MPK6, mitogen-activated protein kinase 6. SND1, secondary wall-associated NAC domain 1. One and two asterisks represent Student’s *t*-test p < 0.05 and 0.01, respectively. Three individual plants were performed as biological replicates for each treatment.

**Fig. S3.**
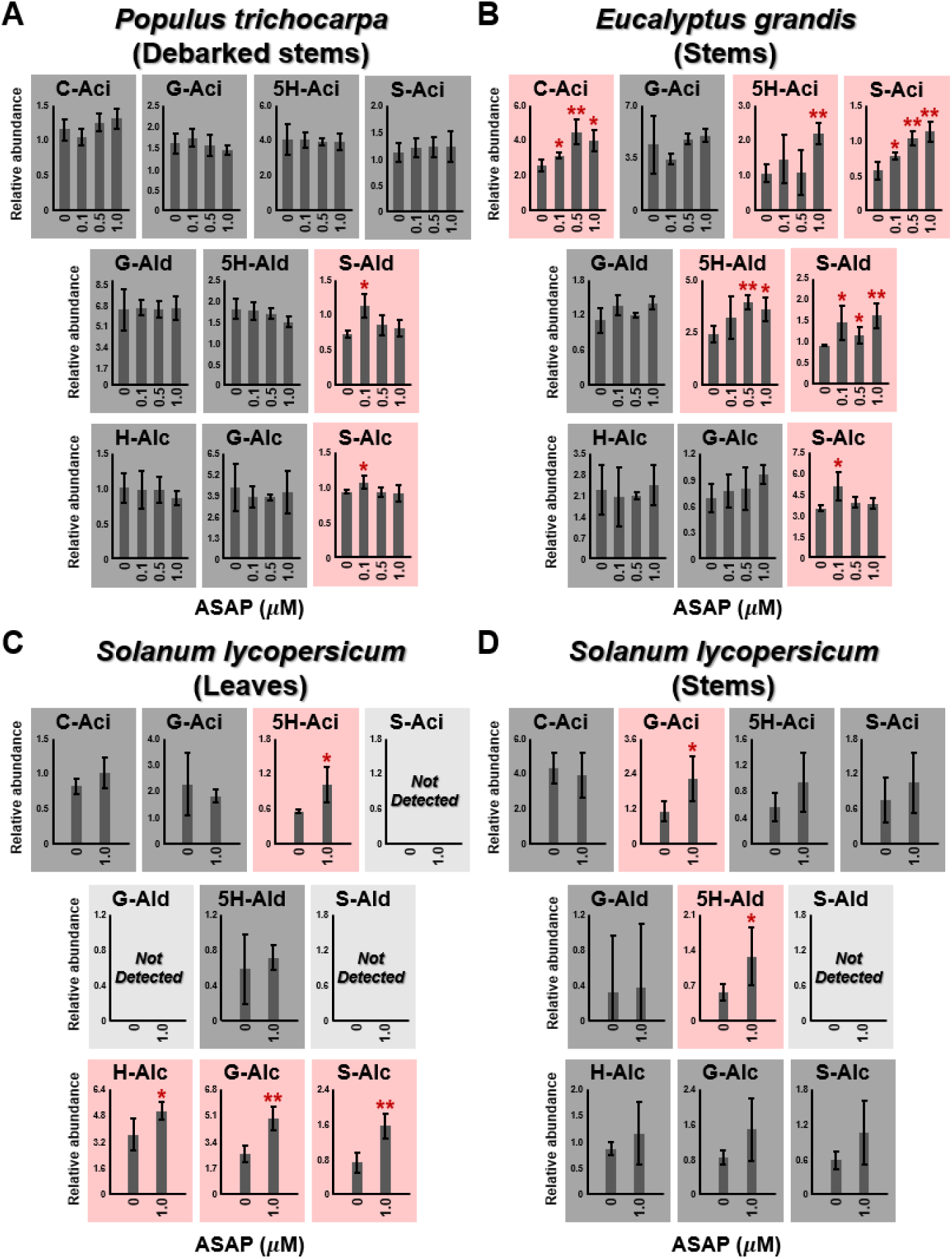
Changes in the monolignol metabolites in three species after ASAP treatment. The changes of monolignol metabolites on (A) the debarked stems of *Populus trichocarpa* and (B) the stems of *Eucalyptus grandis* with different concentrations (0.1 μM, 0.5 μM and 1 μM) of ASAP treatment after one month, and (C) the leaves and (D) the stems of *Solanum lycopersicum* (tomato) with 1 μM ASAP treatment after 24 hours. Ten monolignol metabolites were quantified using LC-SRM-MS analysis. Metabolites in the main fluxes of H-lignin, G-lignin and S-lignin biosynthesis are represented. The detected compounds were highlighted, pink for significant up-regulation, dark grey for no significant change, light grey for no detection. C-Aci, caffeic acid; G-Aci, ferulic acid; 5H-Aci, 5-hydroxyferulic acid; S-Aci, sinapic acid; G-Ald, coniferaldehyde; 5H-Ald, 5-hydroxyconiferaldehyde; S-Ald, sinapaldehyde; H-Alc, 4-coumaryl alcohol; G-Alc, coniferyl alcohol; S-Alc, sinapyl alcohol. One and two asterisks represent Student’s *t*-test p < 0.05 and 0.01, respectively. Three individual plants were performed as biological replicates for each treatment.

**Fig. S4.**
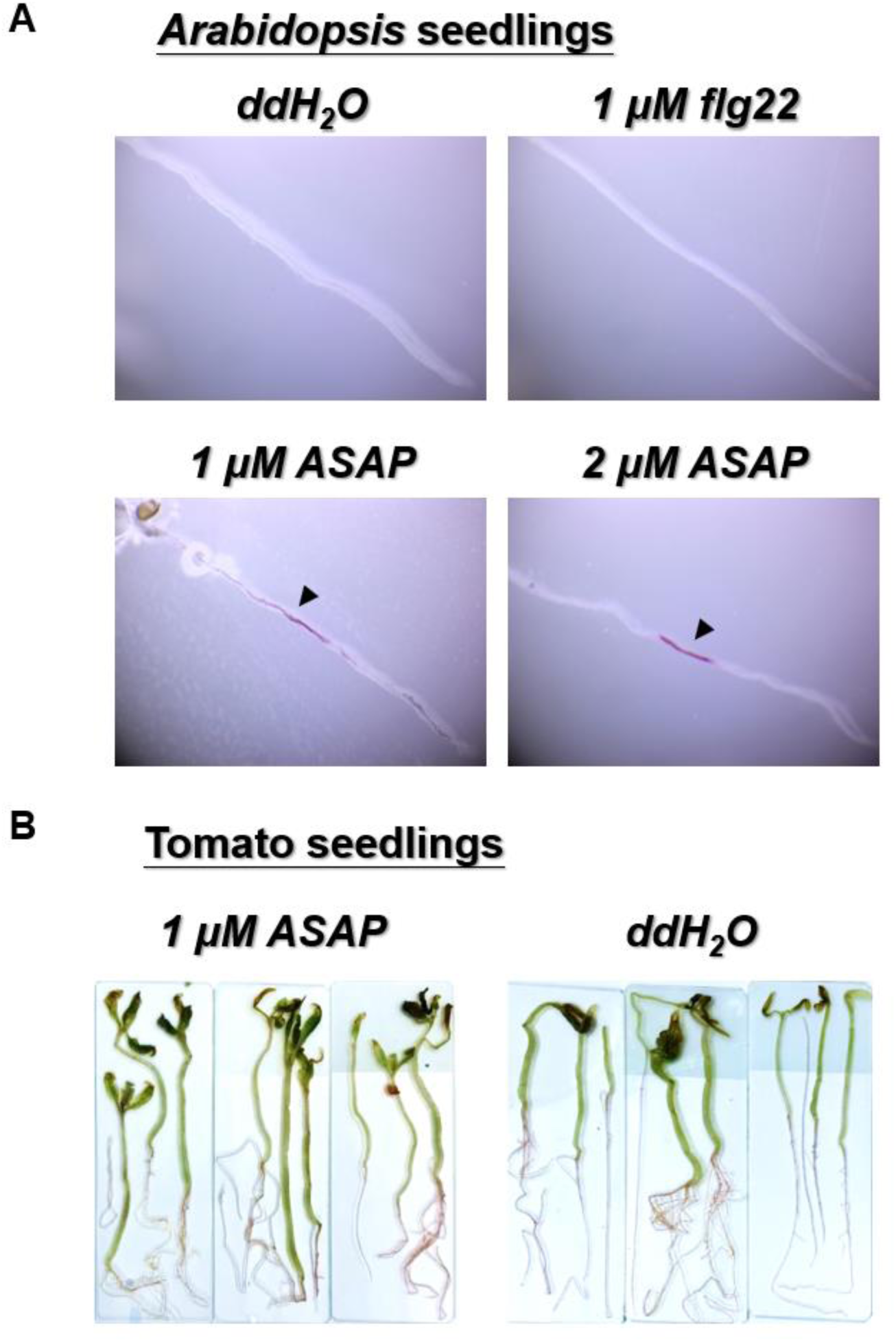
Detection of ASAP-induced lignin deposition in *Arabidopsis* and tomato seedlings. (A) Eight-day-old *Arabidopsis thaliana* seedlings treated with ASAP (1 or 2 µM), flg22 (1 µM) peptides or ddH_2_O (control) for 24 hours. (B) Three-week-old *Solanum lycopersicum* (tomato) seedlings treated with ASAP (1µM) or ddH_2_O (control) for 24 hours. These seedlings were incubated with 40% HCl for 3-5 min, and then moved onto a glass slide, then immersed using 5% phloroglucinol in 95% ethanol for around 15 seconds. The lignified tissues were showed as the pink-red coloration. More than 30 individual plants were performed as biological replicates for each treatment.

**Fig. S5.**
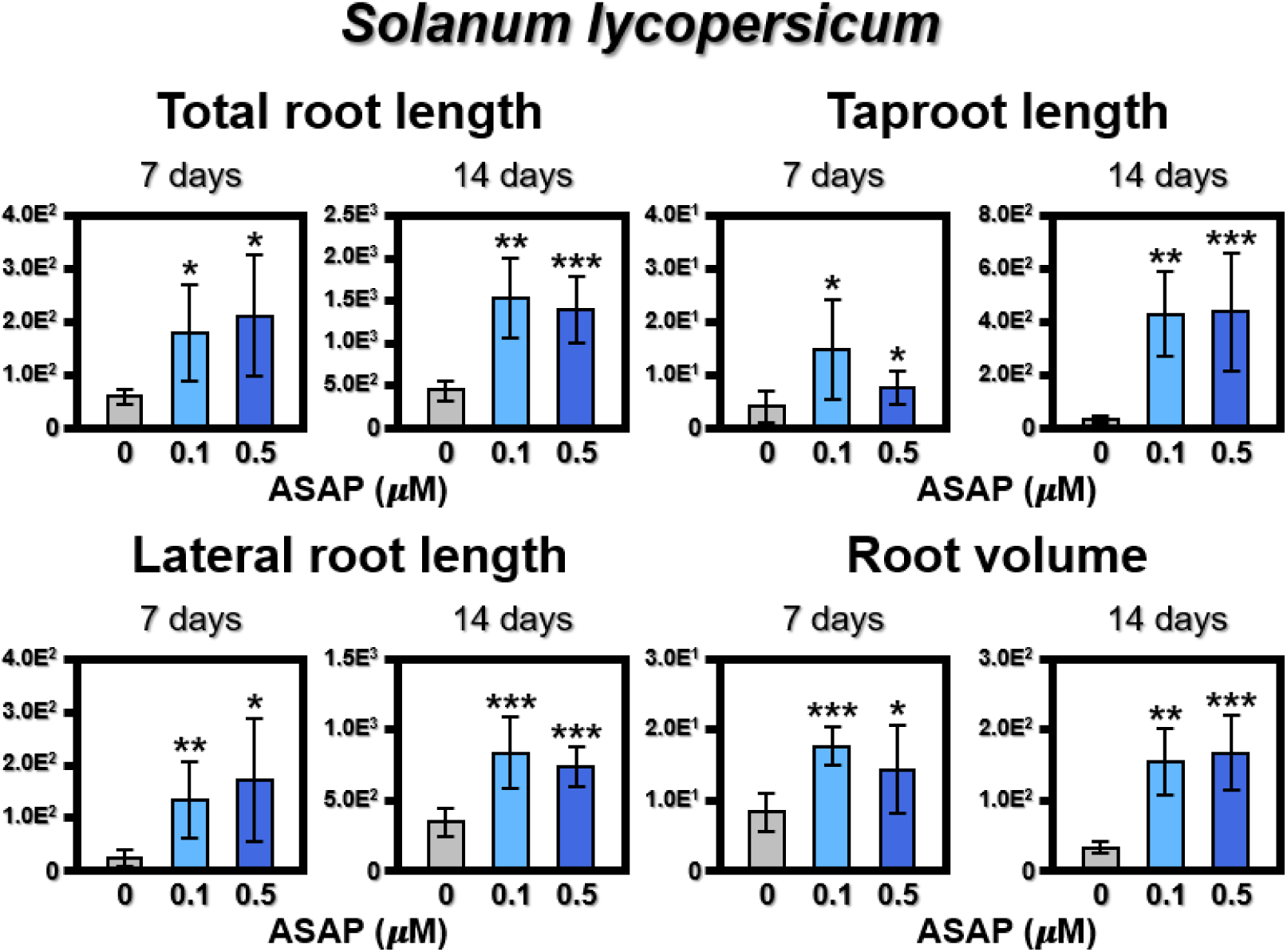
The root development positively regulated by ASAP. Fourteen-day-old *Solanum lycopersicum* seedlings were treated with 0.1 μM or 0.5 μM ASAP for 7 or 14 days. The phenotypes of tomato roots were observed by high-resolution image scanning. Total root length, taproot length, lateral root length, and root volume were observed and calculated. One, two and three asterisks represent Student’s *t*-test p < 0.05, 0.01 and 0.001, respectively. Three individual plants were performed as biological replicates for each treatment.

**Fig. S6.**
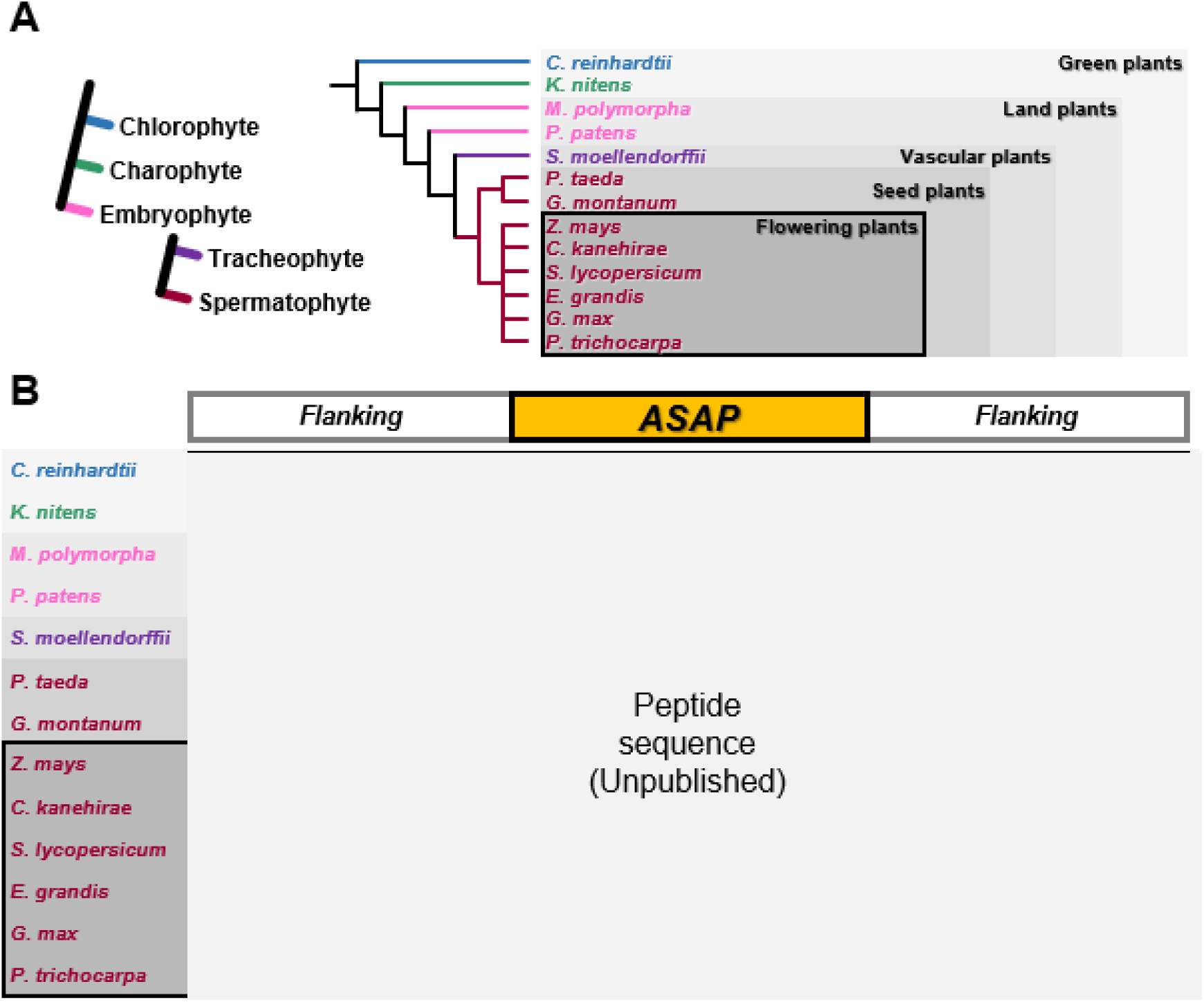
Sequence alignment of ASAP and its homologues across plant kingdom. (A) The phylogeny and the representative species for chlorophyte, charophyte, embryophyte, tracheophyte and spermatophyte. These representative species selected from the different clades of plant kingdom, including *Chlamydomonas reinhardtii*, *Klebsormidium nitens*, *Marchantia polymorpha* and *Physcomitrium patens*, *Selaginella moellendorffii*, *Pinus taeda*, *Gnetum montanum*, *Zea mays*, *Cinnamomum kanehirae*, *Solanum lycopersicum*, *Eucalyptus grandis*, *Glycine max* and *Populus trichocarpa*. (B) The sequences of ASAP with the flanking residues and its homologue sequences in difference species were used to perform the alignment analysis. Yellow represents the identical residues within the ASAP sequence.

**Fig. S7.**
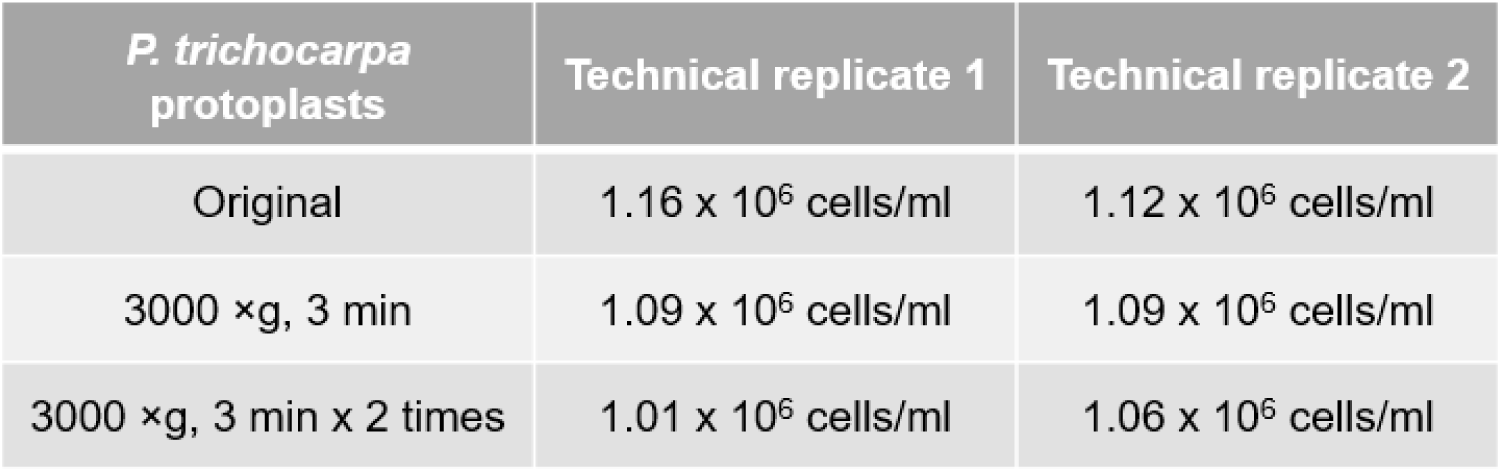
Cell number measurement of the *Populus trichocarpa* protoplasts after centrifugation. Examination of intact cell number before (shown as Original) and after 3000 ×g centrifugation for 3 minutes twice.

**Fig. S8.**
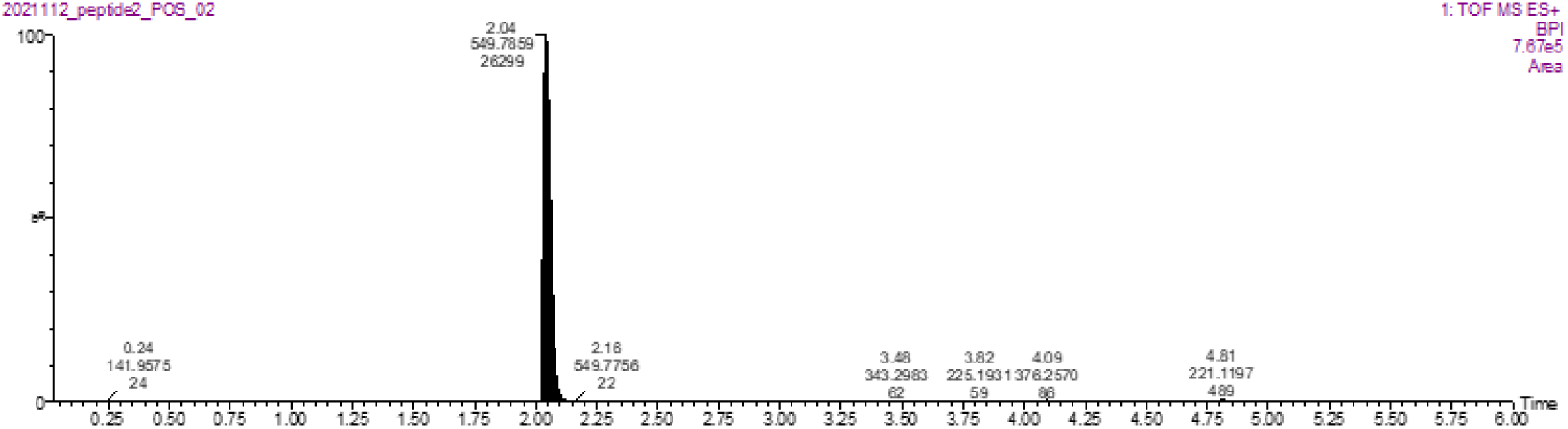
Confirmation of ASAP purity using LC-MS/MS. The mass-to-charge ratio (m/z) of synthetic ASAP sequence (XXXXXXXXXXX) was detected as 549.7859. [M+2H]^2+^: 549.7825. The purity was confirmed to be greater than 95%.

**Fig. S9.**
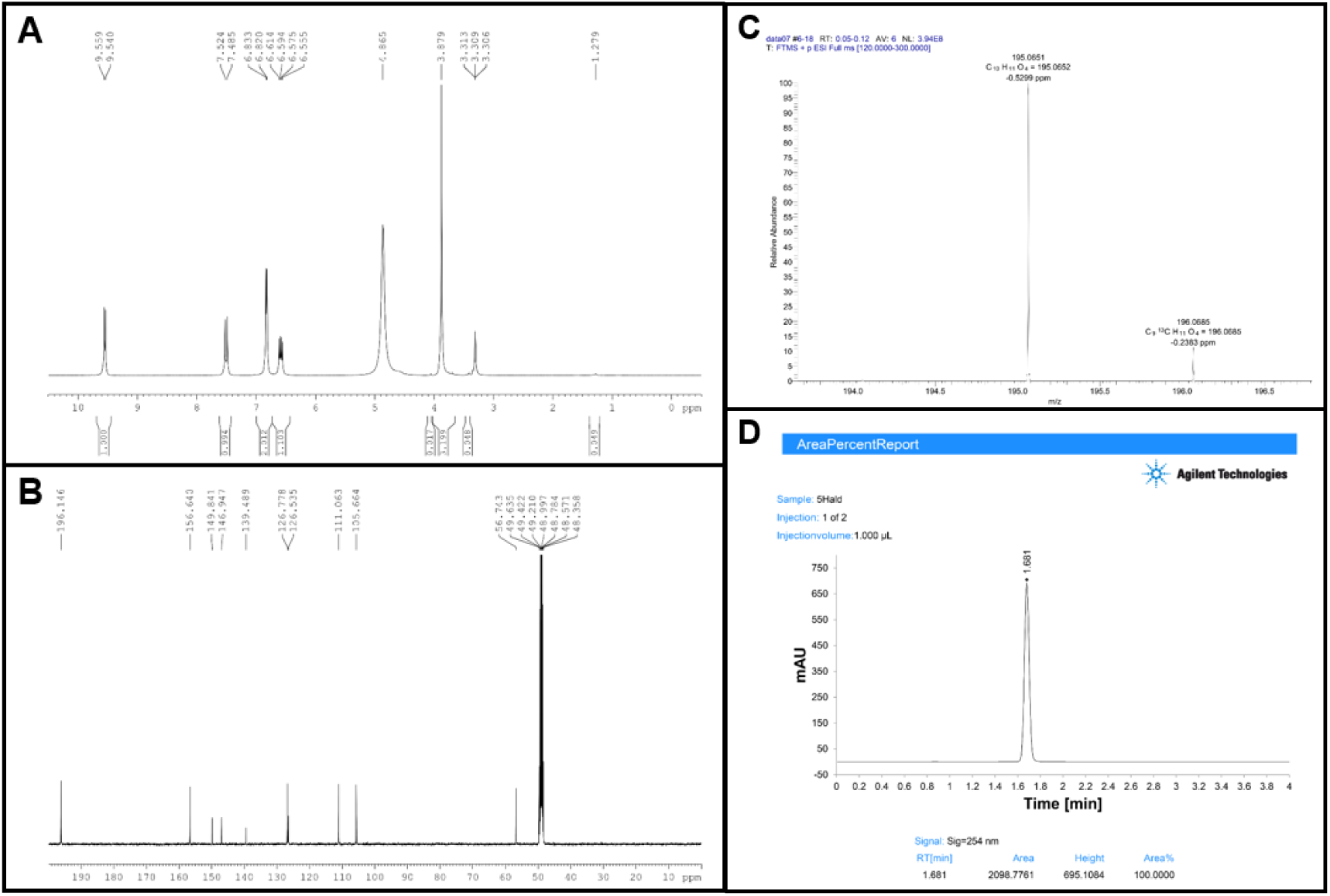
The spectra of synthetic 5-hydroxyconiferaldehyde. (A) ^1^H NMR (400 MHz, MeOD-*d_4_*). ^1^H NMR (400 MHz, MeOD-*d_4_*) δ 9.55 (d, *J* = 8.0 Hz, 1H), 7.50 (d, *J* = 15.6 Hz, 1H), 6.83 (s, 1H), 6.82 (s, 1H), 6.58 (dd, *J* = 15.6, 8.0 Hz, 1H), 3.88 (s, 3H) ppm. (B) ^13^C NMR (100 MHz, MeOD-*d_4_*). ^13^C NMR (100 MHz, MeOD-*d_4_*) δ 196.1, 156.6, 149.8, 146.9, 139.5, 126.8, 126.5, 111.1, 105.7, 56.7 ppm. (C) High-resolution mass spectrum. The formula and m/z were calculated for C_10_H_11_O_4_ [M+H]^+^: 195.0652. Observed m/z: 195.0651. (D) HPLC chromatogram. The mobile phase A is 0.1% formic acid and B is 100% ACN/0.1% FA. The mobile phase consisted of a 75:25 (A:B) isocratic ratio for 4 min. The flow rate was 1.0 ml/min and the injection volume was 1 μl. Peaks were detected at 254 nm.

**Fig. S10.**
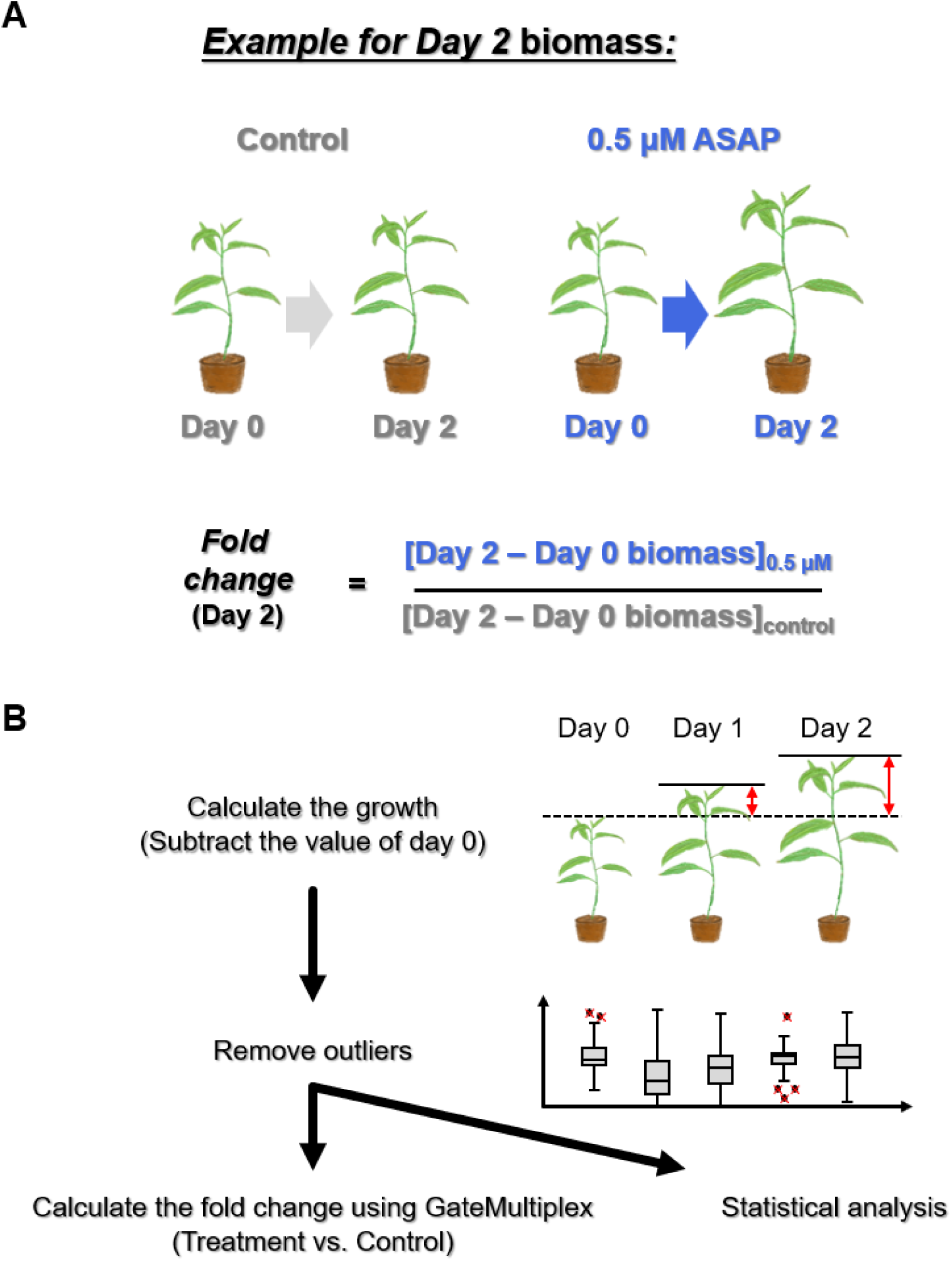
The method for the calculation of changes in digital biomass under ASAP treatment. (A) An example for calculating the fold change of the biomass change. The value of day 0 (the first day for ASAP treatment) biomass was subtracted from the value of the biomass in the following days for each plant. For example, the increased biomass of a plant 2 days upon 0.5 μM ASAP treatment was calculated by subtracting the day 0 biomass from the day 2 biomass after 0.5 μM ASAP treatment, shown as [Day 2 – Day 0 biomass]_0.5μM_. The increased biomass of the control group was also calculated by subtracting the day 0 biomass from day 2 biomass, shown as [Day 2 – Day 0 biomass]_control_, and the fold change of the biomass change after 0.5 μM ASAP treatment for 2 days was calculated by [Day 2 – Day 0 biomass]_0.5μM_ /[Day 2 – Day 0 biomass]_control_. (B) Workflow for the calculation of biomass change. Upon the subtraction of day 0 biomass, the outliers were removed, and the remaining data was applied for the calculation of fold change by GateMultiplex^93^ and the statistical analysis.

## References

1. Hirakawa, Y. & Sawa, S. Diverse function of plant peptide hormones in local signaling and development. Curr. Opin. Plant Biol. 51, 81–87 (2019).

2. Chen, Y. L. et al. Quantitative peptidomics study reveals that a wound-induced peptide from PR-1 regulates immune signaling in tomato. Plant Cell 26, 4135–4148 (2014).

3. Chen, Y. L., Fan, K. T., Hung, S. C. & Chen, Y. R. The role of peptides cleaved from protein precursors in eliciting plant stress reactions. New Phytol. 225, 2267–2282 (2020).

4. Kim, M. J., Jeon, B. W., Oh, E., Seo, P. J. & Kim, J. Peptide signaling during plant reproduction. Trends Plant Sci. 26, 822–835 (2021).

5. Fukuda, H. & Ohashi-Ito, K. Vascular tissue development in plants. Curr. Top. Dev. Biol. 131, 141–160 (2019).

6. Motomitsu, A., Sawa, S. & Ishida, T. Plant peptide hormone signalling. Essays Biochem. 58, 115–131 (2015).

7. Ryan, C. A. & Pearce, G. Polypeptide hormones. Plant Physiol. 125, 65–68 (2001).

8. Franssen, H. J. & Bisseling, T. Peptide signaling in plants. Proc. Natl. Acad. Sci. U.S.A. 98, 12855–12856 (2001).

9. Katsir, L., Davies, K. A., Bergmann, D. C. & Laux, T. Peptide signaling in plant development. Curr. Biol. 21, R356–R364 (2011).

10. Matsubayashi, Y. & Sakagami, Y. Peptide hormones in plants. Annu. Rev. Plant Biol. 57, 649–674 (2006).

11. Tavormina, P., De Coninck, B., Nikonorova, N., De Smet, I. & Cammue, B. P. The plant peptidome: an expanding repertoire of structural features and biological functions. Plant Cell 27, 2095–2118 (2015).

12. Segonzac, C. & Monaghan, J. Modulation of plant innate immune signaling by small peptides. Curr. Opin. Plant Biol. 51, 22–28 (2019).

13. Liu, Z. et al. Phytocytokine signalling reopens stomata in plant immunity and water loss. Nature 605, 332–339 (2022).

14. Ohkubo, Y., Tanaka, M., Tabata, R., Ogawa-Ohnishi, M. & Matsubayashi, Y. Shoot-to-root mobile polypeptides involved in systemic regulation of nitrogen acquisition. Nat. Plants 3, 17029 (2017).

15. Kondo, T. et al. A plant peptide encoded by *CLV3* identified by in situ MALDI-TOF MS analysis. Science 313, 845–848 (2006).

16. Oelkers, K. et al. Bioinformatic analysis of the CLE signaling peptide family. BMC Plant Biol. 8, 1 (2008).

17. Goad, D. M., Zhu, C. & Kellogg, E. A. Comprehensive identification and clustering of CLV3/ESR-related (CLE) genes in plants finds groups with potentially shared function. New Phytol. 216, 605–616 (2017).

18. Mitchum, M. G., Wang, X. & Davis, E. L. Diverse and conserved roles of CLE peptides. Curr. Opin. Plant Biol. 11, 75–81 (2008).

19. De Smet, I., Voss, U., Jürgens, G. & Beeckman, T. Receptor-like kinases shape the plant. Nat. Cell Biol. 11, 1166–1173 (2009).

20. Carella, P., Wilson, D. C., Kempthorne, C. J. & Cameron, R. K. Vascular sap proteomics: providing insight into long-distance signaling during stress. Front. Plant Sci. 7, 651 (2016).

21. Okamoto, S., Suzuki, T., Kawaguchi, M., Higashiyama, T. & Matsubayashi, Y. A comprehensive strategy for identifying long-distance mobile peptides in xylem sap. Plant J. 84, 611–620 (2015).

22. Fukuda, H. & Hardtke, C. S. Peptide signaling pathways in vascular differentiation. Plant Physiol. 182, 1636–1644 (2020).

23. Okamoto, S., Tabata, R. & Matsubayashi, Y. Long-distance peptide signaling essential for nutrient homeostasis in plants. Curr. Opin. Plant Biol. 34, 35–40 (2016).

24. Takahashi, F., Hanada, K., Kondo, T. & Shinozaki, K. Hormone-like peptides and small coding genes in plant stress signaling and development. Curr. Opin. Plant Biol. 51, 88–95 (2019).

25. Ota, R., Ohkubo, Y., Yamashita, Y., Ogawa-Ohnishi, M. & Matsubayashi, Y. Shoot-to-root mobile CEPD-like 2 integrates shoot nitrogen status to systemically regulate nitrate uptake in Arabidopsis. Nat. Commun. 11, 641 (2020).

26. Okamoto, S., Shinohara, H., Mori, T., Matsubayashi, Y. & Kawaguchi, M. Root-derived CLE glycopeptides control nodulation by direct binding to HAR1 receptor kinase. Nat. Commun. 4, 2191 (2013).

27. Takahashi, F. et al. A small peptide modulates stomatal control via abscisic acid in long-distance signalling. Nature 556, 235–238 (2018).

28. Tabata, R. et al. Perception of root-derived peptides by shoot LRR-RKs mediates systemic N-demand signaling. Science 346, 343–346 (2014).

29. Chen, Y. L., Chang, W. H., Lee, C. Y. & Chen, Y. R. An improved scoring method for the identification of endogenous peptides based on the Mascot MS/MS ion search. Analyst 144, 3045–3055 (2019).

30. Villalobos Solis, M. I., et al. A viable new strategy for the discovery of peptide proteolytic cleavage products in plant-microbe interactions. Mol. Plant Microbe Interact. 33, 1177–1188 (2020).

31. Patel, N. et al. Diverse peptide hormones affecting root growth identified in the *Medicago truncatula* secreted peptidome. Mol. Cell. Proteomics 17, 160–174 (2018).

32. Chen, Y. L. et al. XCP1 cleaves Pathogenesis-related protein 1 into CAPE9 for systemic immunity in *Arabidopsis*. Nat. Commun. 14, 4697 (2023).

33. Bi, G. et al. Receptor-like cytoplasmic kinases directly link diverse pattern recognition receptors to the activation of mitogen-activated protein kinase cascades in *Arabidopsis*. Plant Cell 30, 1543–1561 (2018).

34. Asai, T. et al. MAP kinase signalling cascade in *Arabidopsis* innate immunity. Nature 415, 977–983 (2002).

35. Meng, X. et al. A MAPK cascade downstream of ERECTA receptor-like protein kinase regulates *Arabidopsis* inflorescence architecture by promoting localized cell proliferation. Plant Cell 24, 4948–4960 (2012).

36. Liu, S. et al. OsMAPK6, a mitogen-activated protein kinase, influences rice grain size and biomass production. Plant J. 84, 672–681 (2015).

37. Gui, J. et al. Phosphorylation of LTF1, an MYB transcription factor in *Populus*, acts as a sensory switch regulating lignin biosynthesis in wood cells. Mol. Plant 12, 1325–1337 (2019).

38. Xu, R. et al. Control of grain size and weight by the OsMKKK10-OsMKK4-OsMAPK6 signaling pathway in rice. Mol. Plant 11, 860–873 (2018).

39. Guo, T. et al. *GRAIN SIZE AND NUMBER1* negatively regulates the OsMKKK10-OsMKK4-OsMPK6 cascade to coordinate the trade-off between grain number per panicle and grain size in rice. Plant Cell 30, 871–888 (2018).

40. Mao, G. et al. Phosphorylation of a WRKY transcription factor by two pathogen-responsive MAPKs drives phytoalexin biosynthesis in *Arabidopsis*. Plant Cell 23, 1639–1653 (2011).

41. Liu, C., Yu, H., Rao, X., Li, L. & Dixon, R. A. Abscisic acid regulates secondary cell-wall formation and lignin deposition in *Arabidopsis thaliana* through phosphorylation of NST1. Proc. Natl. Acad. Sci. U.S.A. 118, e2010911118 (2021).

42. Zhong, R., Demura, T. & Ye, Z. H. SND1, a NAC domain transcription factor, is a key regulator of secondary wall synthesis in fibers of *Arabidopsis*. Plant Cell 18, 3158–3170 (2006).

43. Jeong, C. Y. et al. Dual role of SND1 facilitates efficient communication between abiotic stress signalling and normal growth in *Arabidopsis*. Sci. Rep. 8, 10114 (2018).

44. Wang, D. et al. The *Arabidopsis* CCCH protein C3H14 contributes to basal defense against *Botrytis cinerea* mainly through the WRKY33-dependent pathway. Plant Cell Environ. 43, 1792–1806 (2020).

45. Zhang, M. & Zhang, S. Mitogen-activated protein kinase cascades in plant signaling. J. Integr. Plant Biol. 64, 301–341 (2022).

46. Haruta, M., Sabat, G., Stecker, K., Minkoff, B. B. & Sussman, M. R. A peptide hormone and its receptor protein kinase regulate plant cell expansion. Science 343, 408–411 (2014).

47. Liu, L. et al. Extracellular pH sensing by plant cell-surface peptide-receptor complexes. Cell 185, 3341–3355 (2022).

48. Wang, J. et al. PEP7 acts as a peptide ligand for the receptor kinase SIRK1 to regulate aquaporin-mediated water influx and lateral root growth. Mol. Plant 15, 1615–1631 (2022).

49. Zhao, Y., Wu, G., Shi, H. & Tang, D. RECEPTOR-LIKE KINASE 902 associates with and phosphorylates BRASSINOSTEROID-SIGNALING KINASE1 to regulate plant immunity. Mol. Plant 12, 59–70 (2019).

50. Yamada, K. et al. The *Arabidopsis* CERK1-associated kinase PBL27 connects chitin perception to MAPK activation. EMBO J. 35, 2468–2483 (2016).

51. Hander, T. et al. Damage on plants activates Ca^2+^-dependent metacaspases for release of immunomodulatory peptides. Science 363, eaar7486 (2019).

52. Dhar, S., Kim, H., Segonzac, C. & Lee, J. Y. The danger-associated peptide PEP1 directs cellular reprogramming in the *Arabidopsis* root vascular system. Mol. Cells 44, 830–842 (2021).

53. Zhang, J., Li, Y., Bao, Q., Wang, H. & Hou, S. Plant elicitor peptide 1 fortifies root cell walls and triggers a systemic root-to-shoot immune signaling in *Arabidopsis*. Plant Signal. Behav. 17, 2034270 (2022).

54. Shen, N., Jing, Y., Tu, G., Fu, A. & Lan, W. Danger-associated peptide regulates root growth by promoting protons extrusion in an AHA2-dependent manner in *Arabidopsis*. Int. J. Mol. Sci. 21, 7963 (2020).

55. Liu, W. & Park, S. W. Underground mystery: Interactions between plant roots and parasitic nematodes. Curr. Plant Biol. 15, 25–29 (2018).

56. Hussain, M. A. & Parveen, G. Determining the damage threshold of root-knot nematode, Meloidogyne arenaria on Vigna unguiculata (L.) Walp. Rhizosphere 27, 100714 (2023).

57. Lu, H. H. et al. Identification of a damage-associated molecular pattern (DAMP) receptor and its cognate peptide ligand in sweet potato (*Ipomoea batatas*). Plant Cell Environ. 46, 2558–2574 (2023).

58. Campbell, L. & Turner, S. R. A comprehensive analysis of RALF proteins in green plants suggests there are two distinct functional groups. Front. Plant Sci. 8, 37 (2017).

59. Huffaker, A. et al. Plant elicitor peptides are conserved signals regulating direct and indirect antiherbivore defense. Proc. Natl. Acad. Sci. U.S.A. 110, 5707–5712 (2013).

60. Huffaker, A., Pearce, G. & Ryan, C. A. An endogenous peptide signal in *Arabidopsis* activates components of the innate immune response. Proc. Natl. Acad. Sci. U.S.A. 103, 10098–10103 (2006).

61. Huffaker, A., Dafoe, N. J. & Schmelz, E. A. ZmPep1, an ortholog of *Arabidopsis* elicitor peptide 1, regulates maize innate immunity and enhances disease resistance. Plant Physiol. 155, 1325–1338 (2011).

62. Yang, R. et al. The SlWRKY6-SlPROPEP-SlPep module confers tomato resistance to *Phytophthora infestans*. Sci. Hortic. 318, 112117 (2023).

63. Lv, W. et al. Comparing the evolutionary conservation between human essential genes, human orthologs of mouse essential genes and human housekeeping genes. Brief. Bioinform. 16, 922–931 (2015).

64. Tang, H. et al. Geometric cues forecast the switch from two- to three-dimensional growth in *Physcomitrella patens*. New Phytol. 225, 1945–1955 (2020).

65. Naramoto, S., Hata, Y., Fujita, T. & Kyozuka, J. The bryophytes *Physcomitrium patens* and *Marchantia polymorpha* as model systems for studying evolutionary cell and developmental biology in plants. Plant Cell 34, 228–246 (2022).

66. Engelen, A. H., Åberg, P., Olsen, J. L., Stam, W. T. & Breeman, A. M. Effects of wave exposure and depth on biomass, density and fertility of the fucoid seaweed *Sargassum polyceratium* (Phaeophyta, Sargassaceae). Eur. J. Phycol. 40, 149–158 (2005).

67. Whitewoods, C. D., et al. *CLAVATA* was a genetic novelty for the morphological innovation of 3D growth in land plants. Curr. Biol. 28, 2365–2376 (2018).

68. Strumillo, M. J. et al. Conserved phosphorylation hotspots in eukaryotic protein domain families. Nat. Commun. 10, 1977 (2019).

69. Canagarajah, B. J., Khokhlatchev, A., Cobb, M. H. & Goldsmith, E. J. Activation mechanism of the MAP kinase ERK2 by dual phosphorylation. Cell 90, 859–869 (1997).

70. Fuglsang, A. T. et al. The binding site for regulatory 14-3-3 protein in plant plasma membrane H^+^-ATPase: involvement of a region promoting phosphorylation-independent interaction in addition to the phosphorylation-dependent C-terminal end. J. Biol. Chem. 278, 42266–42272 (2003).

71. Zhang, W. J., Zhou, Y., Zhang, Y., Su, Y. H. & Xu, T. Protein phosphorylation: A molecular switch in plant signaling. Cell Rep. 42, 112729 (2023).

72. Wang, J. et al. A single transcription factor promotes both yield and immunity in rice. Science 361, 1026–1028 (2018).

73. Ogawa-Ohnishi, M. et al. Peptide ligand-mediated trade-off between plant growth and stress response. Science 378, 175–180 (2022).

74. Ding, S. et al. Phytosulfokine peptide optimizes plant growth and defense via glutamine synthetase GS2 phosphorylation in tomato. EMBO J. 42, e111858 (2023).

75. Stegmann, M. et al. The receptor kinase FER is a RALF-regulated scaffold controlling plant immune signaling. Science 355, 287–289 (2017).

76. Zhang, X., Yang, Z., Wu, D. & Yu, F. RALF–FERONIA signaling: linking plant immune response with cell growth. Plant Commun. 1, 100084 (2020).

77. Abarca, A., Franck, C. M. & Zipfel, C. Family-wide evaluation of RAPID ALKALINIZATION FACTOR peptides. Plant Physiol. 187, 996–1010 (2021).

78. Yoshida, S. “Routine procedures for growing rice plants in culture solution” in Laboratory manual for physiological studies of rice (The International Rice Research Institute, 1976), pp. 61-66.

79. Wang, W., Vignani, R., Scali, M. & Cresti, M. A universal and rapid protocol for protein extraction from recalcitrant plant tissues for proteomic analysis. Electrophoresis 27, 2782–2786 (2006).

80. Tsai, C. F. et al. Sequential phosphoproteomic enrichment through complementary metal-directed immobilized metal ion affinity chromatography. Anal. Chem. 86, 685–693 (2014).

81. Chen, C. W., Tsai, C. F., Lin, M. H., Lin, S. Y. & Hsu, C. C. Suspension trapping-based sample preparation workflow for in-depth plant phosphoproteomics. Anal. Chem. 95, 12232–12239 (2023).

82. Elias, J. E. & Gygi, S. P. Target-decoy search strategy for increased confidence in large-scale protein identifications by mass spectrometry. Nat. Methods 4, 207–214 (2007).

83. Pedrioli, P. G. A. “Trans-proteomic pipeline: a pipeline for proteomic analysis” in Proteome Bioinformatics, vol. 604 of Methods in Molecular BiologyTM (Humana Press, 2010), pp. 213–238.

84. Goodstein, D. M. et al. Phytozome: a comparative platform for green plant genomics. Nucleic Acids Res. 40, D1178–D1186 (2012).

85. Pedrioli, P. G. A. et al. A common open representation of mass spectrometry data and its application to proteomics research. Nat. Biotechnol. 22, 1459–1466 (2004).

86. Holman, J. D., Tabb, D. L. & Mallick, P. Employing ProteoWizard to convert raw mass spectrometry data. Curr. Protoc. Bioinform. 46, 13.24.1–13.24.9 (2014).

87. Chang, W. H. et al. UniQua: a universal signal processor for MS-based qualitative and quantitative proteomics applications. Anal. Chem. 85, 890–897 (2013).

88. Altschul, S. F. et al. Gapped BLAST and PSI-BLAST: a new generation of protein database search programs. Nucleic Acids Res. 25, 3389–3402 (1997).

89. Crooks, G. E., Hon, G., Chandonia, J. M. & Brenner, S. E. WebLogo: a sequence logo generator. Genome Res. 14, 1188–1190 (2004).

90. Thompson, J. D., Higgins, D. G. & Gibson, T. J. CLUSTAL W: improving the sensitivity of progressive multiple sequence alignment through sequence weighting, position-specific gap penalties and weight matrix choice. Nucleic Acids Res. 22, 4673–4680 (1994).

91. Kumar, S., Stecher, G., Li, M., Knyaz, C. & Tamura, K. MEGA X: molecular evolutionary genetics analysis across computing platforms. Mol. Biol. Evol. 35, 1547–1549 (2018).

92. Brien, C., Jewell, N., Watts-Williams, S. J., Garnett, T. & Berger, B. Smoothing and extraction of traits in the growth analysis of noninvasive phenotypic data. Plant Methods 16, 36 (2020).

93. Tsai, N. C. et al. Large-scale data analysis for robotic yeast one-hybrid platforms and multi-disciplinary studies using GateMultiplex. BMC Biol. 19, 214 (2021).

94. Glover, T. J. & Mitchell, K. J. *An Introduction to Biostatistics* (Waveland Press, 2015).

95. Seethepalli, A. et al. RhizoVision Explorer: open-source software for root image analysis and measurement standardization. AoB Plants 13, plab056 (2021).

96. Falk, T. et al. Growing and cultivating the forest genomics database, TreeGenes. Database 2018, bay084 (2018).

97. Perez-Riverol, Y. et al. The PRIDE database resources in 2022: a hub for mass spectrometry-based proteomics evidences. Nucleic Acids Res. 50, D543–D552 (2022).

98. Guo, X., et al. *Chloranthus* genome provides insights into the early diversification of angiosperms. Nat. Commun. 12, 6930 (2021).

